# Chemosensory mechanisms of larval foraging and competitive advantage in the invasive mosquito *Aedes albopictus*

**DOI:** 10.1101/2025.06.07.658027

**Authors:** Dor Perets, Haoqin Ke, Ziyi Yan, Giulia Campli, Evyatar Sar-Shalom, Flavia Krsticevic, Esther Yakir, Robert M. Waterhouse, Yinlang Wang, Philippos A. Papathanos, Jonathan D. Bohbot

**Affiliations:** Department of Entomology, The Hebrew University of Jerusalem, Israel; Agriculture Gene Engineering Research Center of the Ministry of Education, Northeast Normal University, China; SIB Swiss Institute of Bioinformatics, Lausanne, Switzerland; Department of Ecology and Evolution, University of Lausanne, Switzerland

**Keywords:** *Aedes*, odorant receptor, larva, indole, chemosensation

## Abstract

The invasive Asian tiger mosquito, *Aedes albopictus* (Skuse), and yellow fever mosquito, *Aedes aegypti* (L.) are known to compete for resources during the larval stage, often resulting in the ecological displacement of *Ae. aegypti* by *Ae. albopictus*. The chemosensory system plays a pivotal role in larval foraging behavior and may contribute to the competitive advantage. Here, we employed comparative transcriptomics and functional characterization of odorant receptors (ORs) to investigate species-specific differences in larval olfaction. Notably, we uncovered functional variation within the conserved olfactory indole receptor clade, indicating distinct ecological adaptations across species and life stages. We also developed a novel approach to functionally characterize the larval sensory cone and mapped its receptor neuron projections to two key brain regions: the antennal lobe and the subesophageal ganglion. This study provides new insights into the molecular and neural basis of chemosensory-driven behavior in mosquito larvae and highlights the potential role of olfaction in shaping interspecies competition and ecological success.

## Introduction

The yellow-fever mosquito *Aedes aegypti* and the Asian tiger mosquito *Aedes albopictus* are two of the most important invasive mosquito disease vectors of public health concern today [1-4]. These two species evolved in different geographical regions and diverged approximately 35 to 60 million years ago [5, 6]. *Ae. aegypti* originated from the sub-Saharan African forests and is thought to have spread worldwide during the transatlantic slave trafficking and intercontinental shipping of the 15th to 17th centuries [7]. *Ae. albopictus* originates from tropical and temperate Southeast Asia and during the later part of the 20th century spread to other parts of the world via international trade [8, 9]. The spread of *Ae. albopictus* to higher, temperate latitudes has been facilitated by its ability to overwinter, a trait that is absent in *Ae. aegypti* [9, 10, 11]. Due to the rapid spread of *Ae. albopictus* and its vector competency, understanding its biology has become a central focus of public health research and vector control strategies.

When *Ae. aegypti* and Ae albopictus co-occur in the same geographic regions, *Ae. aegypti* is often displaced by *Ae. albopictus*, or the two species segregate into urban and rural habitats, respectively [12-14]. Numerous studies have documented the displacement of *Ae. aegypti* by *Ae. albopictus*, further supporting the competitive advantage of *Ae. albopictus* in shared habitats (**Fig. 1A**) [15-28]. Mechanistic hypotheses for the dominance of *Ae. albopictus* include (i) superiority of *Ae. albopictus* in larval competition [22, 29, 30] (ii) differential susceptibility of *Aedes* species to protozoan parasites, and (iii) interspecific mating favoring *Ae. albopictus* [30-32]. When sympatric, *Ae. albopictus* larvae exhibit greater resilience to low resources than *Ae. aegypti* [30]. A comparative study indicates that *Ae. albopictus* larvae have superior resource-harvesting ability than *Ae. aegypti* [31]. Another study has shown that in insects, a larger chemosensory receptor repertoire is associated with a broader host or habitat range, enabling species to detect and respond to a wider array of environmental cues, which may contribute to their ecological success [33]. Based on these observations, we hypothesized that the larval dominance of *Ae. albopictus* may, in part, be mediated by differences in chemosensory-guided foraging strategies [31].

**Figure 1.**
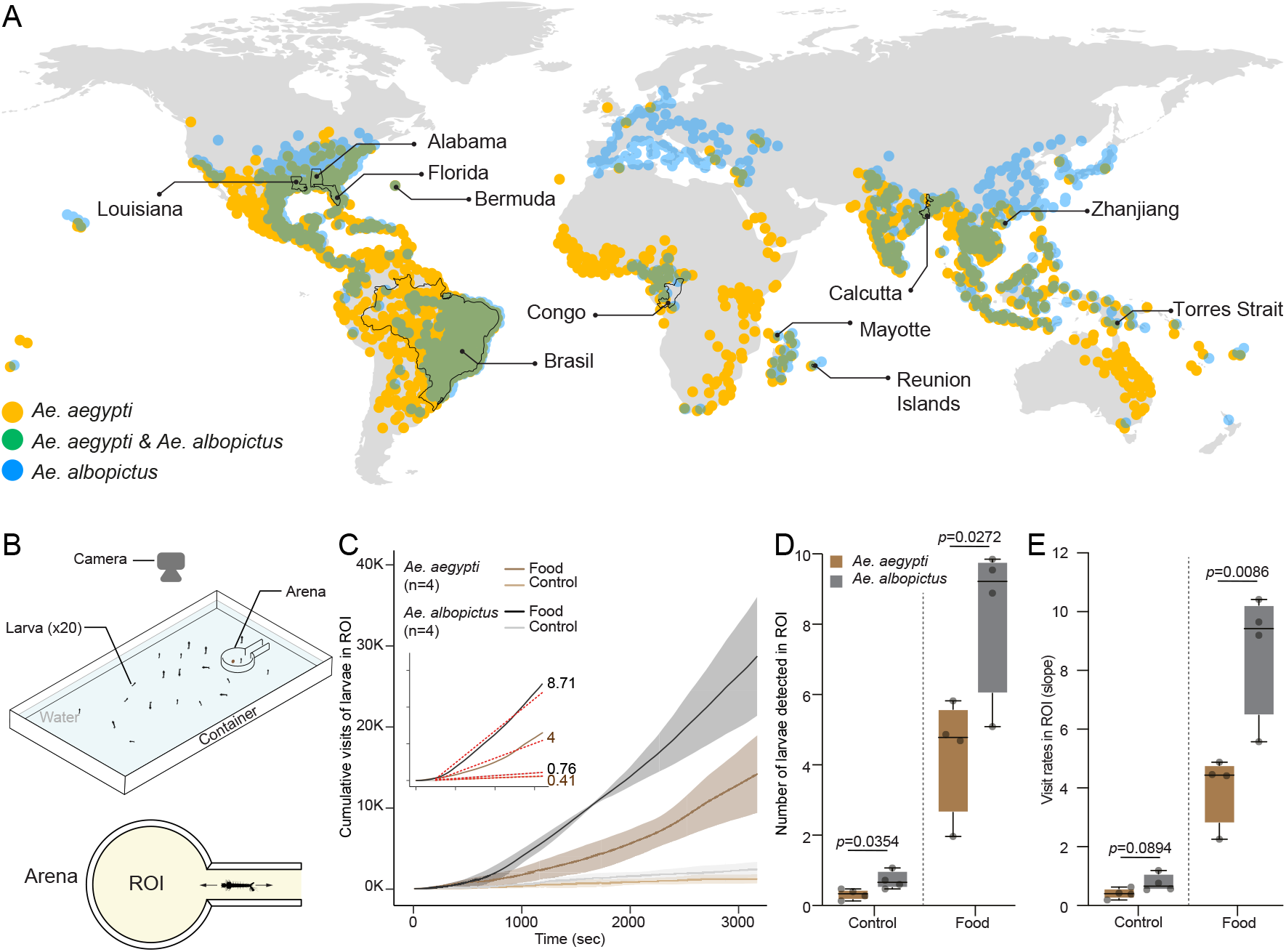
Comparative foraging behavior of *Ae. aegypti* and *Ae. albopictus* larvae in a bioassay. **A**) Reported or hypothetical competitive displacement of *Ae. aegypti* by *Ae. albopictus* in the world. Illustration adapted from Nie & Feng, 2023 [14-28]. **B**) A schematic representation of the experimental setup for the larval bioassay. The setup includes a water-filled container with an arena connected to a corridor on one side. The arena contains a designated region of interest (ROI), marked in the bird’s-eye view at the bottom, where larval movements are quantified. **C**) The cumulative sum of larvae detected in the ROI over a one-hour period for both *Ae. aegypti* and *Ae. albopictus* larvae under food and control conditions. The linear regression trend lines (red dotted lines) illustrate the rates of larval entry into the ROI, with slopes (numbers displayed) indicating the speed of foraging activity. **D**) The average number of larvae detected in the ROI during the one-hour assay for both species under control and food conditions. *Ae. albopictus* larvae displayed a significantly higher average number in the food treatment compared to *Ae. aegypti* larvae (*p* = 0.0272). **E**) The rate of larval entry into the ROI, as quantified by the slopes of the linear regression lines in Panel B. The rate is significantly higher for *Ae. albopictus* in the food treatment (*p* = 0.0086).

The mosquito larval antenna is the main larval chemosensory appendage and is likely used for the detection of food and predators [34]. Compared to the adult, the larval antenna has a significantly lower number of chemosensory organs. It harbors eight sensilla, including one antennal hair, the antennal sensory cone, a basiconic peg organ, a long chaetica sensillum, three long trichoid sensilla, and the sinusoidal peg organ [35]. The large sensory cone on the antenna houses twelve bipolar neurons [35-37]. Additionally, there are three antennal hairs each containing two neurons and three antennal hairs innervated by one neuron each. This anatomical arrangement results in a total of 16 neurons associated with the antennal chemosensory sensilla in the larvae [35, 36], compared to approximately 1600 odorant receptor neurons (ORNs) found in the adult antenna [38]. Axons from the sensory neurons of the antenna project to the larval antennal lobe, a simplified neuropil structure that serves as the primary processing center for olfactory signals [34, 39]. The antennal lobe further projects to higher brain regions, including the deuterocerebrum and the subesophageal ganglion (SOG), which are implicated in the integration of sensory input and motor output for behaviors such as foraging and navigation [39].

The chemosensory gene repertoires of mosquitoes have been the focus of decades of research, but their tissue- and development-specific expression, especially during the larval stage, remains mostly unexplored [40-43]. *Odorant receptor* (*Or*) gene expression in the larval antenna has only been described in *Ae. aegypti* and *Anopheles gambiae*, with a study showing stage-specific expression of some *Ors* [40, 41]. The gene expression profiles of other chemosensory gene families, such as *gustatory receptors* (*Gr*s), *ionotropic receptors* (*Ir*s), *odorant-binding proteins* (Obps), and *pick-pocket* receptors (*Ppks*), have not been examined in the larval antenna. These olfactory proteins play a role in the transduction of chemical signals associated with the identification of food sources, conspecifics, and potential threats [34, 44]. To date, only a small number of *Or*s including *olfactory indolergic receptors* (*IndolOr*s) [45-47] and the 1-octen-3-ol receptor [48], have been shown to be expressed in larvae. In adults, olfactory information from the antenna first transits through the antennal lobe, a structure composed of synaptic glomeruli where axons of olfactory receptor neurons, local neurons, and projection neurons synapse [49]. The existence of such an analogous structure in the larval brain is not well understood [39,49]. The limited knowledge of the chemosensory system in mosquito larvae likely stems from the primary research focus on adult mosquitoes, particularly females, whose olfactory system plays a crucial role in host-seeking and thus disease transmission.

In this study, we performed a comparative investigation of *Ae. aegypti* and *Ae. albopictus* larvae using behavioral assays, antennal olfactory transcriptomics, and functional characterization of IndolORs to evaluate species-specific responses to indolic compounds. Our findings demonstrate that *Ae. albopictus* larvae exhibit significantly higher foraging activity than *Ae. aegypti*, leading us to explore differential chemosensory gene expression as a potential basis for this behavioral divergence. To support mechanistic insights, we established novel single sensillum recording (SSR) protocols, optimized immunohistochemical labeling, and developed anterograde neuronal tracing techniques in *Ae. albopictus*. Neuroanatomical mapping revealed that antennal sensory neurons project to a distinct brain region homologous to the adult antennal lobe, providing a structural framework for future comparative analyses of larval olfactory processing between these ecologically important mosquito species.

## Results

### Larvae of *Ae. albopictus* are more active foragers than *Ae. aegypti*

To compare the relative foraging activities of *Ae. albopictus* to *Ae. aegypti*, we developed a behavioral assay (**Fig. 1A**) to measure the attraction to larval food placed in an arena within the container. Larval visits in the Region of Interest (ROI) within the arena (**Fig1A**) were recorded using a camera and counted throughout each experiment, in the presence of food, or no food as a control. Over the length of the experiment, we examined the temporal dynamics of visits for each species and food treatment (**Fig. 1B, Fig. S1**). The number of *Ae. albopictus* larvae entering the ROI was two-fold higher than *Ae. aegypti* (**Fig. 1C**). In the absence of food, an average of 0.7188 *Ae. albopictus* larvae entered the ROI compared to 0.3177 in *Ae. aegypti* (unpaired *t*-test, t = 2.703, df = 6, *p* < 0.0354) (**Fig. 1C**). The presence of food stimulated visits in the ROI of both species, but *Ae. albopictus* larvae were detected in significantly greater numbers compared to *Ae. aegypti*. The mean number of larvae entering the ROI was 8.341 for *Ae. albopictus*, compared to 4.337 for *Ae. aegypti* (unpaired *t*-test, t = 2.903, df = 6, p < 0.0272) (**Fig. 1C**). To test whether *Ae. albopictus* larvae enter and subsequently explore the ROI faster than *Ae. aegypti*, we fitted our data points to a linear model and calculated the slope of each treatment. The visit rate of both species did not significantly change in the absence of food (unpaired *t*-test, t = 2.024, df = 6, p < 0.0894) (**Fig. 1D**). In the presence of food however, the rate of visits of *Ae. albopictus* was more than double the rate of *Ae. aegypti* larvae (unpaired *t*-test, t = 3.833, df = 6, p < 0.0086).

### *Ae. albopictus* larval antenna expresses a larger *Or* repertoire than *Ae. aegypti*

To begin exploring the genetic basis of the differential foraging activities between these two species, we manually re-annotated the latest *Ae. albopictus* genome assembly (AalbF5) using 439 *Ae. aegypti* chemosensory genes [50] to identify orthologous relationships and construct a phylogenetic tree, resulting in 479 putative chemosensory proteins in *Ae. albopictus* (**Fig. 2A, Fig. S2**). The *Or, Gr, Ir, Obp* and Ppk clades of both species comprised 251, 187, 283, 140 and 82 proteins, respectively. The tree of all *Aedes* chemosensory genes branched as expected according to the five chemosensory gene families (**Fig. 2A**), suggesting that annotations and coding predictions were accurate [51]. We also constructed independent phylogenetic trees for each olfactory gene family (**Fig. S3-7**). We set a threshold of 65% amino-acid identity to determine orthology between proteins of the two species (all data for orthology analysis can be found in https://doi.org/10.5281/zenodo.17292855/Annotations,/Fasta_files, /Orthology_classification, /Similarity_matrices). Each chemosensory gene family exhibited different levels of 1:1 orthologies and species-specific gene expansions (**Fig. 2B**).

**Figure 2.**
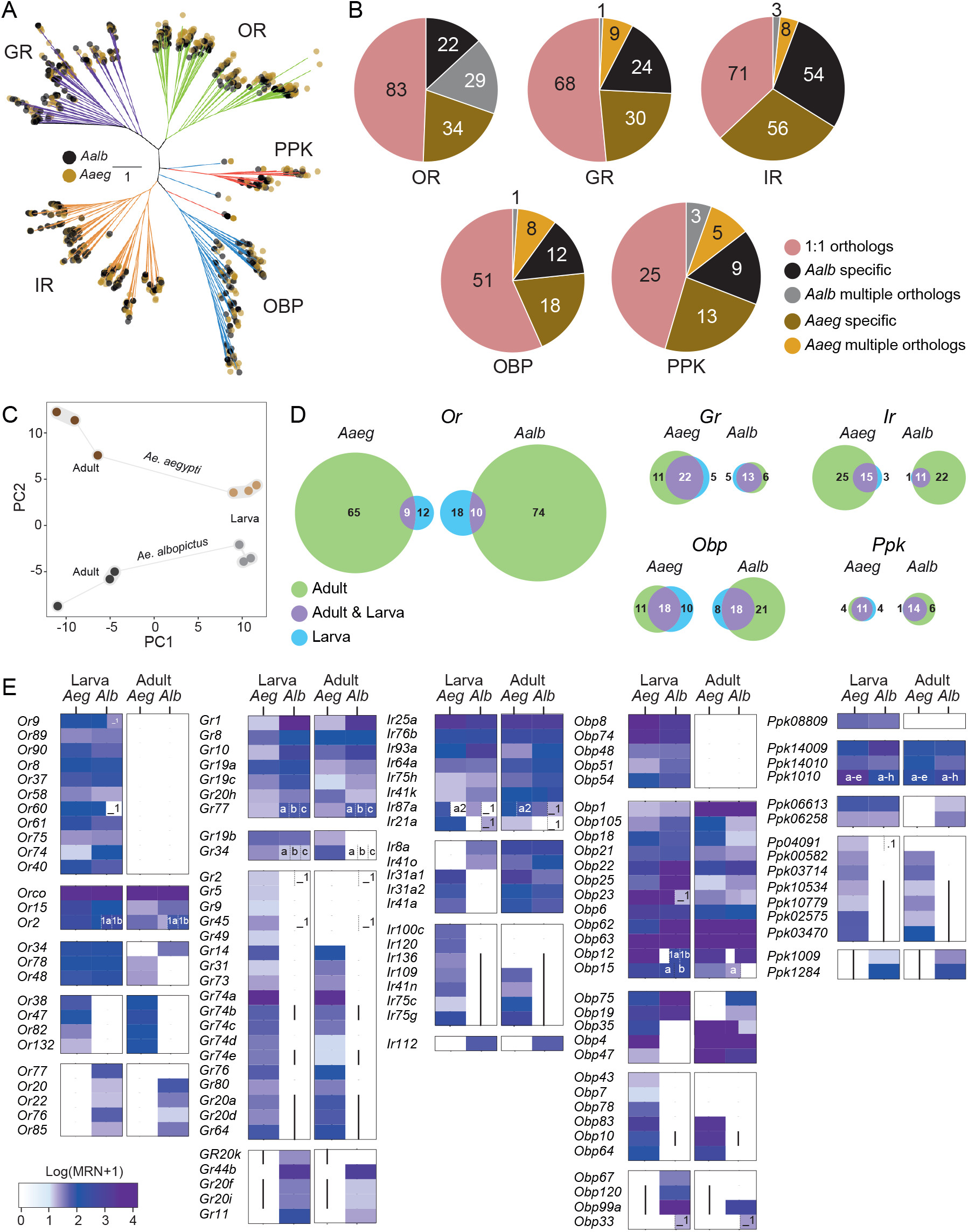
Overview of chemosensory gene profiles in *Ae. aegypti* and *Ae. albopictus*. **A**) An unrooted phylogenetic tree of 439 chemosensory genes from *Ae. aegypti* and 479 chemosensory genes from *Ae. albopictus*. The tree includes genes from the *Or, Gr, Ir, Obp & Ppk* gene families, with branches color-coded by gene family. **B**) Pie charts showing the distribution of chemosensory genes across *Or, Gr, Ir, Obp & Ppk* families in *Ae. aegypti* and *Ae. albopictus*. Conserved 1:1 orthologs are shown in light red, while species-specific expansions are shown in brown (*Ae. aegypti*) and black (*Ae. albopictus*). Cases where multiple proteins from one species shared >65% amino acid identity with a single ortholog in the other species are considered multiple orthologs and represented in light brown (*Ae. aegypti*) and grey (*Ae. albopictus*). **C**) Principal component analysis (PCA) of chemosensory gene expression (*e*.*g*., *Or, Gr, Ir, Obp & Ppk*) in the antennae of *Ae. aegypti* and *Ae. albopictus* at both larval and adult stages. Each point represents a biological replicate, with axes representing principal components PC1 and PC2. **D**) Venn diagrams displaying the overlap and distribution of stage-specific (green and light blue) and developmentally-shared (purple) chemosensory genes between larvae and adults in both species. The threshold for expression was set at mean ratio normalized counts (MRN) **≥** 10. Separate Venn diagrams are provided for *Or, Gr, Ir, Obp & Ppk* gene families. **E**) Heatmaps showing the expression levels (log2(Mean reads normalised + 1)) of chemosensory genes across the five main chemosensory gene families in the larval and adult antennae of *Ae. aegypti* (*Aeg*) and *Ae. albopictus* (*Alb*).

Next, we dissected antennae from L3-L4 larvae and male and female adults from both species and sequenced three replicates of each using Illumina NovaSeq X. RNAseq reads were mapped to the AalbF5 *Ae. albopictus* and L5.2 *Ae. aegypti* genome assemblies. Principal component analysis of the expression of chemosensory genes clustered replicates closer than samples spanning different developmental stages or species, as expected, and larval antennae were more similar to each other than adult antennae between species (**Fig. 2C**). In both species, the larval antenna expressed distinct subsets of *Ors* while *Gr, Ir, Obp* and Ppk genes showed greater overlap between developmental stages (**Fig. 2D**).

Overall, the total number of expressed chemosensory genes in the *Ae. albopictus* larval antenna was similar to that of *Ae. aegypti*. We observed an increase in the number of expressed *Ors* and *Ppks* (**Fig. 2D**). *Ae. albopictus* larvae expressed seven more *OR*s in the larval antenna than *Ae. aegypti* (**Fig. 2D**), of which two are duplications of *Or2, Or9* in *Ae. albopictus* (**Fig. 2E)**. Of the 12 and 18 *OR*s that are specifically expressed in the larval in *Ae. aegypti* and *Ae. albopictus*, respectively, 12 are 1:1 orthologs (*Or9, 89, 90, 8, 37, 58, 60, 61, 75, 74* and *40*). In terms of expression levels in the larval antennae, besides *Orco* that is highly expressed, the most highly expressed *ORs* were *Or2, Or9* and *Or8* and *Or15*. Among the remaining *Ors* expressed in the larval antennae, their expression in the adult antenna was variable and expressed in one or both species. *Gustatory receptor* gene expression levels were lowest among all chemosensory genes (**Fig. 2E**). Interestingly, the number of *GR*s expressed in the larval antenna of *Ae. aegypti* was approximately 33% higher than in *Ae. albopictus*. Several *GR*s were commonly expressed in both species, including subunits of the CO_2_ receptor complex (*Gr1*) [52], as well as *Gr19a/c, Gr8, Gr10, Gr20h, Gr77*, and *Gr34* (**Fig. 2E, Fig. S9**). Homologs of the *D. melanogaster* sugar *Gr5* and *9* receptors [53] were expressed in the *Ae. aegypti*, but not in the *Ae. albopictus* larval antenna. By comparison, the sugar *D. melanogaster Gr11* homolog was expressed in the *Ae. albopictus* larval antenna [54]. The mosquito ortholog (*Gr34*) of the *D. melanogaster* fructose receptor *Gr43a* was expressed in the larval antennae of both species. Of the three Ir co-receptors only *Ir25a* and *Ir76b* were expressed in the larval antenna, while the *Ir8a* co-receptor was expressed only in *Ae. albopictus* larvae [55]. Only three or one ligand sensing *Ir* subunits were specifically expressed in the larvae antenna, the remainder of the *Irs* were shared between larvae and adults (**Fig. 2D,E**). Twenty-eight and twenty-six *Obp* genes were expressed in the larval antenna of *Ae. aegypti* and *Ae. albopictus*, respectively, with approximately half of them being larval-specific (**Fig. 2D,E**). *Odorant-binding protein* gene expression was either species or stage-specific.

*Pickpocket* gene expression was largely shared between stages, with four *Ppk*s exhibiting larva-specific expression in *Ae. aegypti*, while *Ae. albopictus* larvae showed stage-specific expression of only a single gene, *Ppk04091*, in the antenna. We performed differential expression analysis between adult and larval antennae and quantified the number of differentially expressed genes (DEGs) in each species. In both *Ae. aegypti* and *Ae. albopictus*, the number of chemosensory genes upregulated in the larval antenna exceeded those upregulated in the adult antenna (**Fig. 3A**). Given the central role of the antenna as a sensory organ across life stages, we next quantified and categorized stage-biased chemoreceptors in each species (**Fig. 3B**). This analysis revealed a conserved pattern in the number of differentially expressed chemosensory genes between species. Notably, the DEG analysis identified stage-specific or stage-biased expression of multiple chemosensory genes in each species (**Fig. 3C, Fig. S8–S13**).

**Figure 3.**
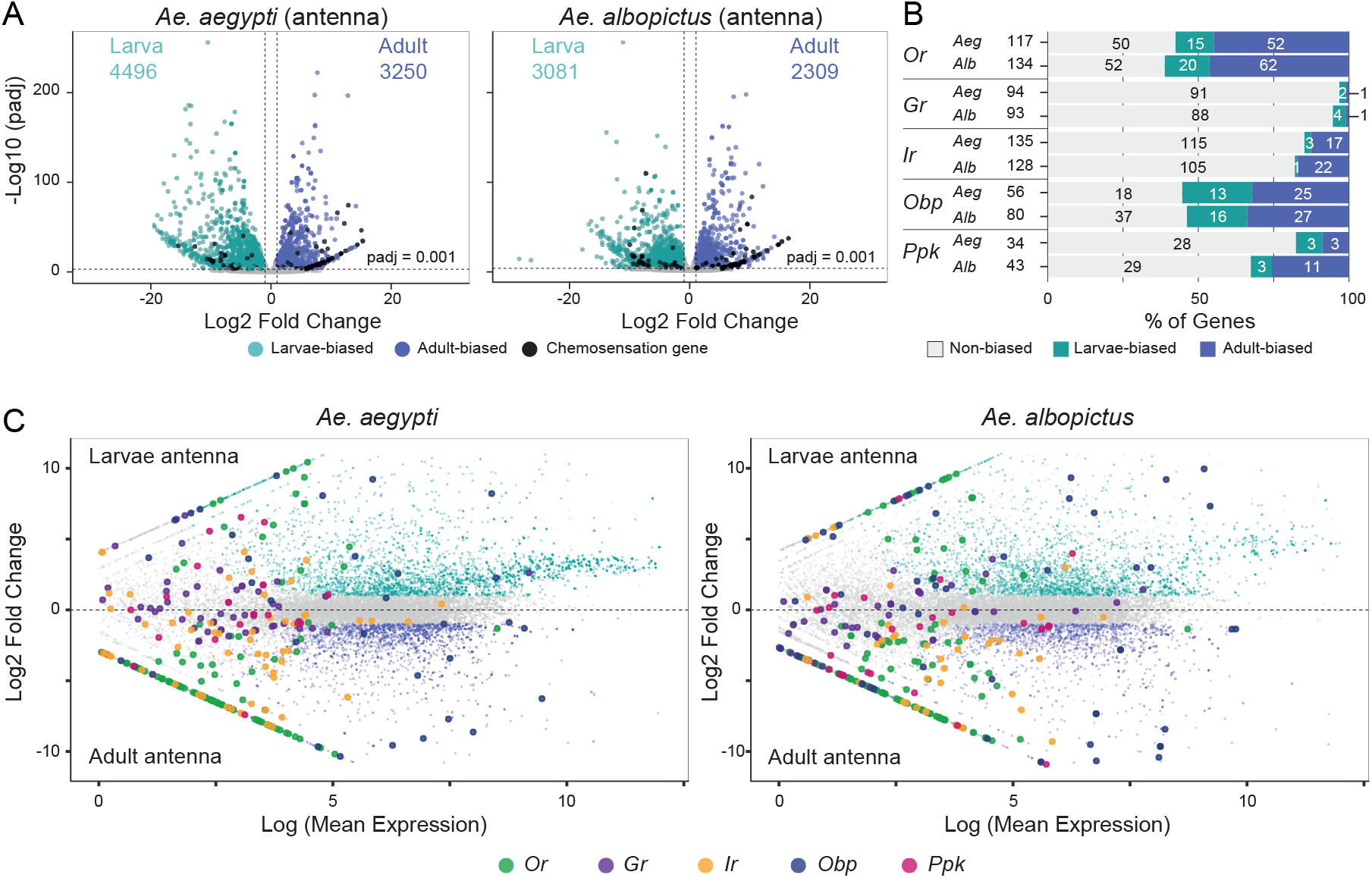
Overview differential expression analysis of larval versus adult antenna in *Ae. aegypti* and *Ae. albopictus*. **A**) Volcano plot of larval antenna versus adult antenna in each species. Thresholds for differentially expressed genes are P adjusted value (Padj) <=0.001 and Log2FoldChange > 1. Larvae-biased genes are in turquoise and adult-biased genes are in blue, black dots are chemosensory genes (*e*.*g*., *OR, GR, IR, OBP, PPK*). Numbers at the top of each side of the plots indicate total DEGs in each stage. **B**) Quantification of differentially expressed chemosensory genes across the *Or, Gr, Ir, Obp & Ppk* families in both species. **<u>C</u>**) Scatter plots of larval antenna versus adult antenna in *Ae. aegypti* and *Ae. albopictus*. Genes with significant stage differential expression (Padj < 0.001) are shown in turquoise or blue for larva/adult biased respectively, while non-significant genes between the stages are depicted in gray.

### IndolORs function in *Ae. albopictus* larvae

Transcriptomic and phylogenetic analyses confirmed that indolORs in *Ae. aegypti* and *Ae. albopictus* share conserved amino acid sequences and antennal expression patterns. Given that the odorant tuning properties of these receptors are well-characterized in *Ae. aegypti*, we focused our efforts on the functional characterization of indolORs in *Ae. albopictus*. Given that *Ae. albopictus* and *Ae. aegypti* compete at the larval stage, which may be mediated by the detection of bacteria-derived organic compounds, we compared the response of indolORs to indole and skatole. Using quantitative RT-PCR, we confirmed that in *Ae. albopictus, Orco* expression was significantly higher in adult antennae compared to larvae (**Fig. 4A**). This difference was more pronounced in *Ae. aegypti*. We also found that *Or2* expression was detected in both larval and adult antennae, *Or9* was larval-specific, and *Or10* was adult-specific (**Fig. 4A**; **Fig. S13**). A highly similar developmental expression pattern was observed in *Ae. aegypti* with *Or2, Or9*, and *Or10*. The expression pattern in *Ae. aegypti* was consistent with previous reports [40, 45, 47]. Full statistical analyses of the qPCR experiments are provided in the Zenodo qPCR folder (https://doi.org/10.5281/zenodo.17292855).

**Figure 4.**
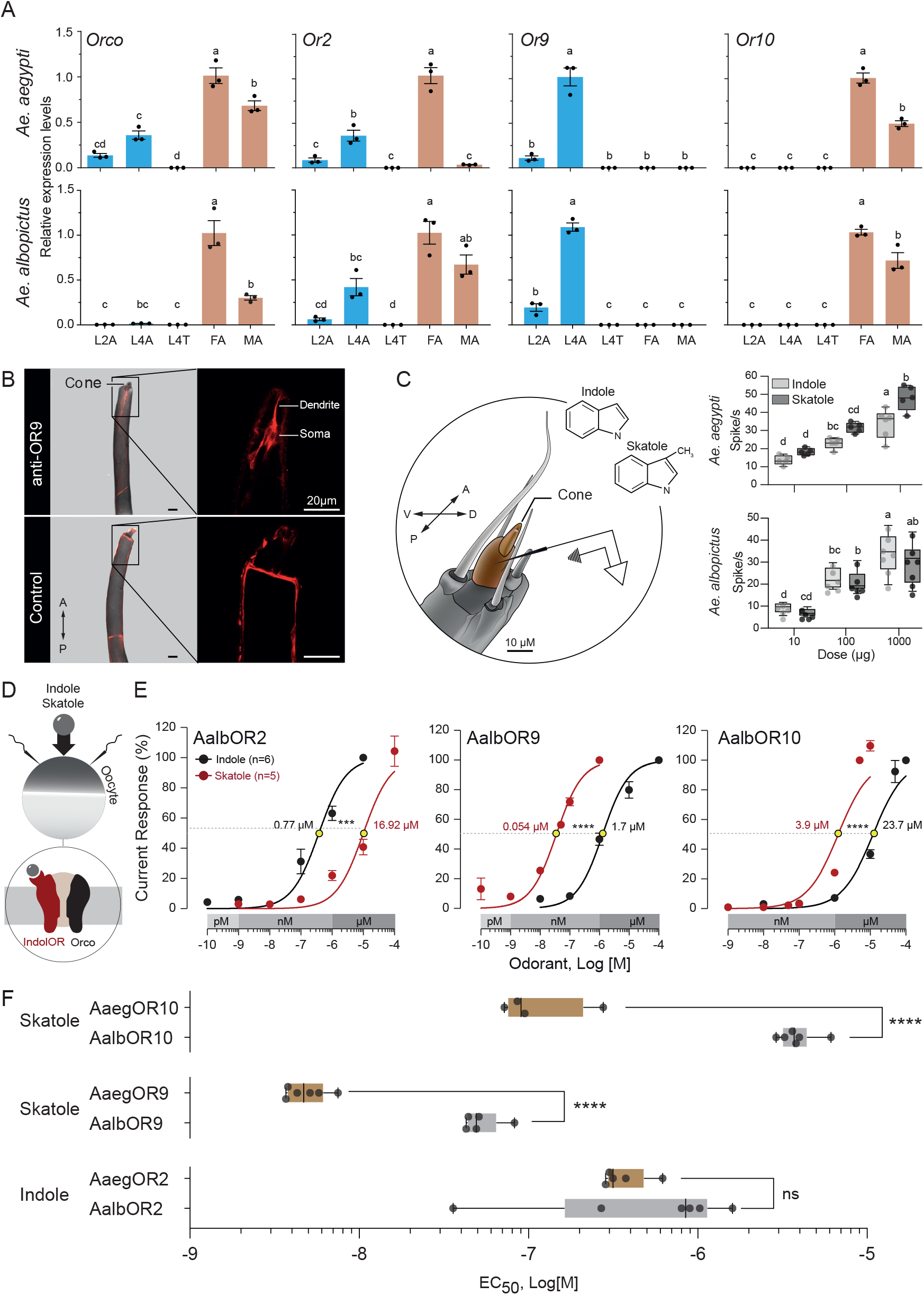
Expression and functional characterization of odorant receptors in *Ae. Aegypti* and *Ae. albopictus*. **A**) RT qPCR *indolOR* expression levels of *OrcO, Or2, Or9*, and *Or10* in *Ae. aegypi* and *Ae. albopictus* across developmental stages: second instar larvae (L2A), fourth instar larvae (L4A), larval terminal segments (L4T), female adults (FA), and male adults (MA). Expression levels are normalized, and letters (a, b, c) indicate statistical differences (Ordinary one-way ANOVA, Tukey multiple comparisons tests) between groups (*p* < 0.05). **B**) Immunostaining of *Ae. albopictus* larval antennae using anti-OR9 antibodies (top) and control staining (bottom). The labeled cone structure in the larval antenna shows localization of OR9 in the dendrites and soma. Scale bars indicate magnifications. **C**) Left: Schematic representation of the *Aedes* larval antenna, highlighting the cone structure where responses to indole and skatole were measured. Right: Dose-dependent SSR electrophysiological responses (spikes) to indole and skatole at varying doses (10 µg, 100 µg, and 1000 µg) for *Ae. aegypti* (top) and *Ae. albopictus* (bottom). Boxplots represent the median, interquartile range, and individual data points for each dose. Statistical tests for *Ae. aegypti* responses (ordinary one-way ANOVA, F(5,25) = 31.89, *p* < 0.0001; Tukey’s multiple comparison test, df = 25; Indole 10 vs Indole 100 Δ = –9.4, *p* < 0.0001; Indole 100 vs Indole 1000 Δ = –10.83, *p* = 0.0143; Skatole 10 vs Skatole 100 Δ = –13.4, *p* = 0.0003; Skatole 100 vs Skatole 1000 Δ = –16.2, *p* < 0.0001; Indole 10 vs Skatole 10 Δ = –4.6, *p* = 0.2072; Indole 100 vs Skatole 100 Δ = –8.6, *p* = 0.0982; Indole 1000 vs Skatole 1000 Δ = –13.97, *p* = 0.0011). Statistical tests for *Ae. aegypti* responses (ordinary one-way ANOVA, F(5,30) = 15.57, p < 0.0001; Tukey’s multiple comparison test, df = 30; Indole 10 vs Indole 100 Δ = –13.4, *p* = 0.0222; Indole 100 vs Indole 1000 Δ = –11.43, *p* = 0.0396; Skatole 10 vs Skatole 100 Δ = –13.93, *p* = 0.0159; Skatole 100 vs Skatole 1000 Δ = –8.24, p = 0.2392; Indole 10 vs Skatole 10 Δ = 2.2, *p* = 0.9944; Indole 100 vs Skatole 100 Δ = 1.67, *p* = 0.9977; Indole 1000 vs Skatole 1000 Δ = 4.86, *p* = 0.7322). **D**) Schematic of the experimental setup for heterologous expression of *Ae. albopictus Ors* with *Orco* in *Xenopus* oocytes. The setup measures receptor responses to indole and skatole. **E**) Dose-response curves of AalbOR2, AalbOR9, and AalbOR10 to indole (black) and skatole (red). The x-axis shows the logarithmic concentration of the ligands (Molar), and the y-axis represents the normalized current response (%). EC_50_ values are marked with yellow circles. Unpaired parametric t-test results (for full results see the main text) between responses to the ligands are indicated (\*\*\**p* < 0.001, \*\*\*\**p* < 0.0001). **F**) Comparison of EC_50_ values (log M) for indole and skatole across orthologous odorant receptors in *Ae. aegypti* (*Aaeg*) and *Ae. albopictus* (*Aalb*). One-way ANOVA statistical tests results between the species (\*\*\*\**p* < 0.0001; ns: not significant). Boxplots show the median, interquartile range, and individual replicates. The EC_50_ values for the *Ae. aegypti* indolORs are from previous studies [45,47].

Staining the larval antenna using an *OR9* antibody localized the receptor protein to the dendrite and soma of neurons in the sensory cone (**Fig. 4B**). Using the base recording method (**Fig. S14A**), we observed that 1-octen-3-ol, indole, and skatole elicited increased spike frequencies from larval antennal cones in both *Ae. albopictus* and *Ae. aegypti*. However, we were unable to distinguish between different neuron types in our recordings (**Fig. S14B**). Both *Ae. aegypti* and *Ae. albopictus* larvae exhibited dose-dependent increases in neuronal activity in response to indole and skatole (**Fig. 4C**) (ordinary one-way ANOVA followed by Tukey’s multiple comparison test, see Fig. 4C for detailed statistical analyses). In *Ae. aegypti*, responses to skatole were higher than to indole at the highest dose.

To quantify the sensitivity of IndolOR proteins from *Ae. albopictus*, we expressed AalbOR2, OR9, and OR10 in *Xenopus* oocytes (**Fig. 4D**) and recorded their current responses to increasing concentrations (100 pM to 10 µM) of skatole and indole (**Fig. 4E, S15**). AalbOR2 displayed an EC_50_ of 0.77 µM and 10.09 µM for indole and skatole, respectively (unpaired t-test: *p* = 0.0005, t = 5.249, df = 9). AalbOR9 exhibited low nanomolar sensitivity to skatole (EC_50_= 0.054 µM), and low micromolar sensitivity to indole (EC_50_ = 1.7 µM, unpaired t-test: *p* < 0.0001, t = 12.50, df = 8). The adult-specific AalbOR10 exhibited lower sensitivity to skatole (EC_50_=3.9 µM), and lower sensitivity to indole (EC_50_ = 23.7 µM, unpaired t-test: *p* < 0.0001, t = 12.48, df = 8) than skatole (**Fig. 4E, S15**). To evaluate the functional conservation and sensitivity of IndolORs between *Ae. aegypti* and *Ae. albopictus*, we compared their EC_50_ values to their cognate ligand, skatole or indole (**Fig. 4F**). For skatole, AalbOR10 and AalbOR9 exhibited a significantly lower EC_50_ value than AaegOR10 (One-way ANOVA: *p* < 0.0001). There was no significant difference in sensitivity between AalbOR2 and AaegOR2 for indole (One-way ANOVA: *p* = 0.9863).

### The larval antennal nerve projects to brain regions consistent with the antennal lobe and the suboesophageal ganglion

To visualize the projection patterns of chemosensory neurons of the larval antenna to the brain, we used anterograde dye staining in *Ae. aegypti* and *Ae. albopictus* (**Fig. 5B-C**). General brain staining highlighted structures typically described in insects, including a large supraesophageal ganglion (SuEG), two optic lobes (OLs), and the subesophageal ganglion (SOG) (**Fig. 5B-C**). The SuEG and the SOG are connected by two circumesophageal connectives (CCs) surrounding the oesophagous foramen (OF). The antennal nerve mostly projected ipsilaterally to a region (**Fig. 5D-E**) previously associated with the antennal lobe of *Ae. aegypti* larvae [39]. These projections were always observed on the same side of the brain where the antenna was stained (**Fig. 5, Fig. S17-18**). In this area the antennal nerve showed multiple globular tangles reminiscent of glomeruli (**Fig. 5D-E, Fig. S17-18**). Immediately below the candidate antennal lobe (AL), the antennal nerve descended ventrally into the CC with additional globular tangles consistent with the antennal mechanosensory and motor centers (AMMC) and proceeded to the SOG where multiple arborizations were visualized (**Fig. 5F-G**). Notably, we observed in 2 of the 3 preparations contralateral projections in the SOG (**Fig. 5F-G, Fig. S17-18**).

**Figure 5.**
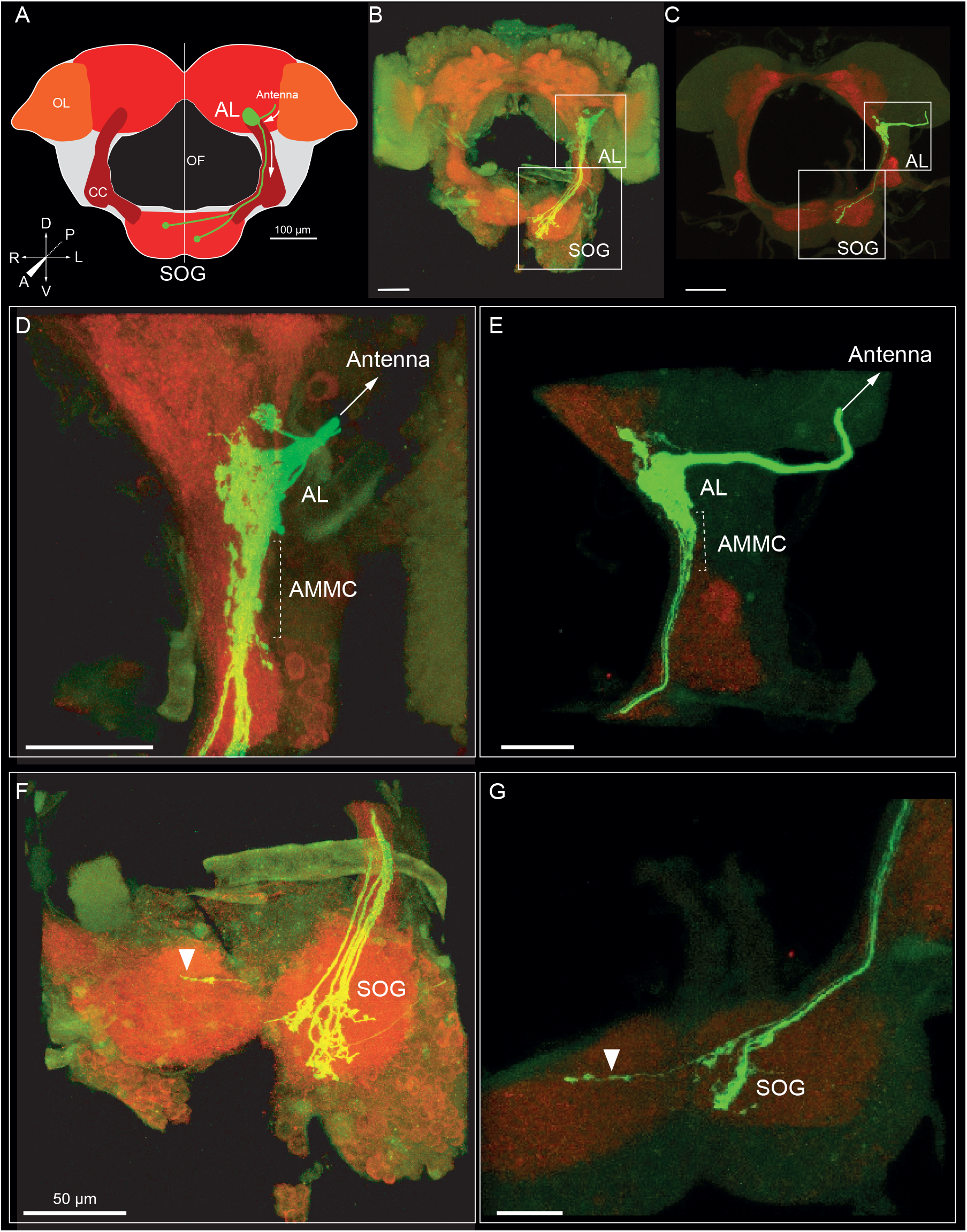
Anterograde staining of the whole larval antenna highlighting the neural projections from the antenna to the antennal lobe (AL) and the suboesophageal ganglion (SOG). **A**) Schematic of the larval brain (adapted from [39]). Optic lobe (OL), deutocerebrum (DE), mushroom bodies (MB), circumesophageal connective (CC), and oesophageous foramen (OF). **B-C**) Bright field image of an L4 larva whose antenna is covered with a capillary for anterograde tracing. *Ae. albopictus* in **B** and *Ae. aegypti* in **C**,**D**,**E**) Magnified region of the projections to the presumed AL and antennal mechanosensory and motor center (AMMC*). Ae. albopictus* in **D** and *Ae. aegypti* in **E. F-G**) Magnified region of the ipsilateral and contralateral (white triangle) projections to the SOG. *Ae. albopictus* in **F** and *Ae. aegypti* in **G**.

## Discussion

Our study provides the first comparative description of the olfactome profiles of two *Aedes* mosquito larval antennae. The superior foraging activity observed in *Ae. albopictus* could be driven by differences in navigation behaviors, foraging strategies, or chemosensation. Both *Aedes* species have been shown to share similar navigation behaviors and foraging strategies [56, 57]. In the absence of food, we found no significant foraging differences between the species. However, when food was present, *Ae. albopictus* larvae displayed a faster exploration rate than its competitor. These findings suggest that chemosensory detection is the primary factor driving these behavioral differences.

To begin exploring the potential genetic basis of chemosensory detection differences between the two species, we generated an RNA-seq atlas of larval and adult antennae and carefully re-annotated the latest, highly-repetitive chromosome-level *Ae. albopictus* genome assembly. Overall, the *Ae. albopictus* genome assembly contains a similar number of chemosensory genes of *Ae. aegypti*.

We observed differences between these mosquito species in the expression of different chemosensory genes in the larval antennae that could form the basis for the future discovery of key genes mediating the superiority of *Ae. albopictus* in larval habitats. Interestingly, we found that expression of orthologous *Ors* was highly conserved in levels and tissue-specificity, but this was not the case for *Grs*, where each species has a unique set of receptors in the larval antenna. In *Ae. aegypti*, the larval antennae expressed receptors homologous to *Drosophila melanogaster* sugar receptors [53], may suggest distinct food preferences compared to *Ae. albopictus. Ae. albopictus* larvae expressed a greater number of *Or*s in the antenna compared to *Ae. aegypti*, with seven additional *Or*s detected. While many receptors were shared between species, several *Or*s were uniquely expressed in one species but not the other. Notably, most of these species-specific *Or*s were expressed across both larval and adult stages, rather than being restricted to a particular developmental stage. These patterns suggest that species-specific differences in olfactory receptor expression are not limited to stage-specific deployment but may reflect broader divergence in chemosensory gene regulation.

In contrast, the larval antenna of *Ae. aegypti* expresses a broader repertoire of *Gr*s, with many receptors detected exclusively at the larval stage. Nevertheless, several *Ae. albopictus*-specific *Gr*s were identified in the larval antenna, including *Gr11*, which is known to play a role in oviposition site selection in adult mosquitoes [54]. The presence of candidate sugar-sensing receptors in larvae raises the question of the potential source of sugar that larvae could find in their aquatic pools. Moreover, among the ionotropic receptors, *Ir8a* was expressed in the larval antenna exclusively in *Ae. albopictus*, but not in *Ae. aegypti*. In *Ae. aegypti, Ir8a* expression is restricted to the adult antenna, the primary chemosensory organ, whereas the two other ionotropic co-receptors, *Ir25a* and *Ir76b*, are also expressed in the maxillary palps and tarsi [55]. All three ionotropic co-receptors (*Ir8a, Ir25a*, and *Ir76b*) were detected in both larval and adult antennae of *Ae. albopictus*, suggesting broader stage-specific deployment in this species.

Previous work in *Ae. aegypti* has shown that *indolOrs*, two of which are expressed in the larval antenna, exhibit heightened sensitivity to indolic microbial volatile organic compounds [45, 46, 47, 57]. We have shown here that both species express *IndolOr2* and *Or9* in their larval antennae. While OR2 of each species responds similarly to indole, AaegOR9 is 10 times more sensitive to skatole than AalbOR9. The low-micromolar EC_50_ value of AalbOR9 suggests either the existence of a more potent, yet unidentified, indole derivative or that this receptor has a broader tuning spectrum than its *Ae. aegypti* homolog. A similar pattern was observed for both OR10s. The heightened sensitivity of *Aaeg*OR9 and the antennal cone to skatole suggests a potential causal relationship, which can be directly tested once mutant lines become available.

The presence of the OR9 protein in the antennal cone suggests that this sensory structure functions as an olfactory organ. Furthermore, this study is among the first to provide a detailed characterization of chemosensory projections from the larval antenna to the brain, providing insights into how sensory signals are processed and integrated in mosquito larvae. To achieve this, we applied single sensillum recording and anterograde staining to investigate the sensory cone and higher-order signal integration in the brain, respectively. This methodological advancement paves the way for future research comparing chemosensory-driven foraging behaviors in these two mosquito species, shedding light on how larval sensory processing contributes to their ecological success.

## Materials and Methods

### Mosquito rearing

Laboratory colonies were maintained in environmental insect chambers under controlled conditions (26 °C, 80% RH, 12:12 h light:dark cycle). Larvae were reared in plastic pans containing 1 L of water and fed a ground mixture of Novocrabs feed (JBL GmbH & Co., Germany) until pupation. Pupae were transferred to plastic cages, and emerging adults were allowed to mate and fed ad libitum on a 10% sucrose solution. Females were provided cattle blood using the HemoTek feeding system. The Foshan-Pavia (FPA) strain of *Ae. albopictus* was obtained from the Bonizzoni laboratory, University of Pavia, Italy, and the Liverpool (LVP) strain of *Ae. aegypti* originated from Prof. Joel Margalit’s laboratory, Ben-Gurion University. While these laboratory strains are widely used in mosquito research, their long-term maintenance under laboratory conditions may limit their representativeness of natural populations.

### Larval foraging behavior

A 30 x 18 cm glass arena was filled with one liter of 24°C water into which a disposable, 3D-printed stimulant maze (“arena”) was placed always with the corridor against the incubator wall in different directions (left/right) in order to avoid positional effects (arena dimensions and experiment layout in **Fig. 1A**). Twenty L4 larvae, that had previously been starved for 2 hours, were used in each experiment and placed in fresh 24°C DDW in each repetition. All experiments were conducted around 13:00 under the same conditions of the rearing. The treatments were: control (empty arena), food tablet (Novocrabs feed: JBL GmbH & Co, Germany). Water was replaced for each new experiment. The larvae were tracked with IR-sensitive camera (Pimoroni MLX90640) for 1h. The data from each treatment were processed manually by counting the number of larvae in ROI for a frame every second (3600∼ frames). Statistical significance between species was determined using parametric unpaired t-test. All raw videos of larva and count matrices of larva inside the ROI are included in the Larva_behaviour in https://doi.org/10.5281/zenodo.17292855 (Aaeg_videos, Aalb_videos, Larva_behaviour_Count_matrices folders).

### Chemosensation genes annotation in AalbF5

Annotation of chemosensation gene orthologs in the *Ae. albopictus* genome assembly was performed by *de novo* gene prediction combined with orthology delineation. The longest isoform for each protein of interest from the *A. aegypti* proteome (AaegL5.0, GCF_002204515.2) was selected, resulting in 390 query proteins (*OR,GR,IR & OBP* from Aaeg_meta_ID file in in https://doi.org/10.5281/zenodo.17292855 **/** Annotations folder). Queries were used for gene annotation in the *Ae. albopictus* genome assembly (AalbF5, GCF_035046485.1) by the gene predictor MetaEuk [58] with default parameters and the – exhaustive-search option. At least one statistically significant prediction was found for 99% of the queries (https://doi.org/10.5281/zenodo.17292855 **/** Annotations folder). Orthology delineation was performed using OrthoLoger v3.0.2 [59] from 19 highly complete Diptera genome annotation sets available from the United States National Center for Biotechnology Information (with Benchmarking Universal Single-Copy Orthologs-BUSCO-Complete Scores for the Diptera lineage dataset of at least of 95%) and the *Ae. albopictus* annotated gene set https://doi.org/10.5281/zenodo.17292855 Annotations/Predictions/7160_and_predictions.fa), including the most significant prediction resulted after each query (https://doi.org/10.5281/zenodo.17292855 **/** Annotations/ Predictions/ AalbF5_best_predictions_all_genes.gff). All proteins from *Ae. aegypti* and *Ae. albopictus* (either from the public annotation or the newly predicted gene annotations) included in the same OrthoLoger-defined orthologous group were considered co-orthologs. Following the automated prediction and orthology delineation, we manually reviewed the top predictions and extracted newly identified *Ae. albopictus* sequences absent from prior annotations. These curated sequences were then included in the subsequent phylogenetic analyses.

### Phylogenetic analysis

Chemosensation genes have been extensively characterized through genomics and transcriptomics studies in *Ae. aegypti* [42, 50]. For finding *Ae. albopictus* orthologs in AalbF5 assembly, 479 sequences obtained from the AalbF5 genome and aligned using MAFFT G-INS-1 alignment strategy [60,61]. Alignment files were used for construction of Maximum-likelihood phylogenetic trees using IQ-TREE [62]. The genes were classified to the five main chemosensory gene-families (*e*.*g. OR, GR, IR, OBP, PPK*) and each gene was given a name by homology to their *Ae. aegypti* orthologs. In cases when an *Ae. albopictus* had more than 1 ortholog to *Ae. aegypti* the second/third ortholog was marked with “_1”/ “_2” (*e*.*g. OR2_1*). The underscore suffix referred to a secondary ortholog which was found in different loci. Cases when a gene has more than 1 alternative transcript, we added a suffix of a letter (*e*.*g OR10a, OR10b*). Percentages of amino acid identity were computed using the MAFFT G-INS-1 alignment and the results were annotated manually on the bootstrap values of each gene family phylogenetic tree (**Figures S4-8**). All data are available at supplementary data: https://doi.org/10.5281/zenodo.17292855 (Annotations, Fasta_files, Orthology_classification, Similarity_matrices).

### Antennal collections and RNAseq analysis

Larvae RNA libraries consisted of 200 larval antennae dissected from L3 - L4 larva, 6dpe Adult RNA libraries consisted of 75 male and 75 female adult antennae. RNA extracted using TRIzol reagent (Invitrogen™ 15596026) and Direct-zol™ RNA MiniPrep (ZYMO research™ R2050). Libraries sequenced at Novogene facility in Singapore using Illumina NovaSeq X, each library sequenced depth of 30M reads 150 bp paired-end reads (Fastq RNA libraries available at NCBI SRA: PRJNA1234527, PRJNA1268156 for *Ae. albopictus* and *Ae. aegypti* respectively). Low quality reads were discarded using Trimmomatic [63] by equal value cutoff of 28. RNA reads were aligned to the AalbF5 *Ae. albopictus* and L5.2 *Ae. aegypti* genome assemblies using HiSAT [64] using the -k 10 parameter, which was adjusted to allow each read to be assigned to up to 10 alignments. Gene count matrices were extracted using FeatureCounts using default settings with ambiguous (-a) read counting. This approach ensures a comprehensive count of multi-mapping reads, which is particularly important for accurately quantifying gene expression in regions with overlapping or closely related transcripts [65]. Raw gene expression tables are available in supplementary data (https://doi.org/10.5281/zenodo.17292855 /expression matrices folder). Differential expression analysis was conducted using DEseq2 to compare the expression patterns of different developmental stages and species using default parameters [66]. Normalization of raw count data was conducted using the DESeq2 median ratio normalization (MRN) method, which is preferred for its ability to effectively correct for compositional biases and differences in sequencing depth across samples, ensuring accurate comparison of gene expression levels. Moreover Fragments Per Kilobase of transcript per Million mapped reads (FPKM) values were calculated using DESeq2. To define gene expression, a cutoff of 10 normalized counts was applied, with a gene considered expressed if its average count across three biological replicates from larval antennae exceeded this threshold. All expression data are available in https://doi.org/10.5281/zenodo.17292855 /expression matrices folder.

### Quantitative real-time PCR

Total RNA was extracted by using TRIzol reagent (Invitrogen, Carlsbad, CA, USA) following the manufacturer. After extraction, the concentration and purity of RNA were checked by a NanoDrop 2000 spectrophotometer (Thermo Fisher Scientific, Waltham, MA, USA). Corresponding cDNAs were synthesized by 0.5 µg total RNA with the First-Strand cDNA Synthesis Kit (Transgen Biotech, Beijing, China). The *β-actin* (LOC5574526, LOC115253654 for *Ae. aegypti* and *Ae. albopictus* respectively) gene was used as an internal control. The specific primers were designed by using Primer 3 (https://bioinfo.ut.ee/primer3-0.4.0/) and qPCR were performed on a LightCycler 480 II Detection System (Roche, Basel, Switzerland) and TransStar Tip Top Green qPCR Supermix kit (Transgen Biotech, Beijing, China). The qPCR conditions were: 94 °C for 30 s, followed by 45 cycles of 94 °C for 5 s, 55 °C for 15 s, and 72 °C for 10 s. Gene expression levels were calculated by using the 2^−ΔΔCT^ method. All qPCRs were performed with at least three technical and three biological replicates. Primers and data used in qPCR found in https://doi.org/10.5281/zenodo.17292855/ qPCR folder.

### Functional characterization of the *Ae. albopictus* indolOR

The *Xenopus laevis* females (pigmented *Xenopus laevis*) were obtained from Nasco (Fort Atkinson, WI, USA). The frogs were reared in a XenoPlus amphibia housing system (Tenciplast, West Chester, PA, USA) under controlled water conditions (pH = 7.6, Temp = 18.2°C, conductivity = 1100 μS). The frogs were fed three times a week with an adult *Xenopus* irradiated diet (Zeigler, Gardners, PA, USA, Prod. No. 316518-18-2412). Whole-cell currents were monitored and recorded using the two-electrode voltage clamp (TEVC) technique. Oocytes were injected with 27.6 nL of RNA (*AalbOR2*-XM_029872179.2, *AalbOR9* - XM_029863963.1, *AalbOR10a* - XM_062854695.1) using the Nanoliter 2010 injector (World Precision Instruments, Inc., Sarasota, FL, USA). Each receptor was co-injected with its obligatory co-receptor *AalbORco* - XM_029877254.2. Injected oocytes were then incubated at 18oC in 24 well-plates containing 1 ml incubation medium as described above for 3 days prior to recording. *OR2,9,10* responses was recorded by measuring whole cell currents using the two-electrode voltage clamp (TEVC) method. Holding potential was maintained at −80 mV using an OC-725C oocyte clamp (Warner Instruments, LLC, Hamden, CT, USA). Oocytes were placed in a RC-3Z oocyte recording chamber (Warner Instruments, LLC, Hamden, CT, USA) and exposed for 8-second-long stimuli. Currents were allowed to return to baseline between odorant applications.

Data acquisition was carried out with a Digidata 1550A and pCLAMP10 (Molecular Devices, Sunny-vale, CA, USA). Holding potential was maintained at −80 mV using an OC725C oocyte clamp (Warner Instruments, LLC, Hamden, CT, USA). Oocytes were placed in a RC-3Z oocyte recording chamber (Warner Instruments, LLC, Hamden, CT, USA) and exposed to three second-long stimuli. All compounds were solubilized in 200 μL of DMSO prior to dilutions in ND96 buffer supplemented with 0.8 mM CaCl. Data acquisition was carried out with a Digidata 1550A and pCLAMP10 (Molecular Devices, Sunnyvale, CA, USA). TEVC raw recording traces and data tables are in https://doi.org/10.5281/zenodo.17292855 **/** Pharmacology folder.

### Single-sensillum recording of *Ae. albopictus* larval antennae

Indole (99%, Sigma-Aldrich) and skatole (98%, Sigma-Aldrich) were firstly diluted in dimethyl sulfoxide (DMSO) to make a 10% (m/v) stock solution, and then diluted to the desired concentrations before use. The 4th instar larvae were fixed into a 10 μL plastic pipette tip, and the head of the larva was exposed and fixed by dental wax. The exposed antennae were adhered to a coverslip with double-sided tape under a microscope (LEICA Z16 APO, Germany), the reference electrode (sharpened tungsten wire) was inserted into the eye, and the recording electrode was inserted into the middle of sensory cone of larva’s antenna until a stable spontaneous potential with a high signal-to-noise ratio was achieved. For the stimulus, 10 μL of solution was added on a 1 cm × 2.5 cm filter paper and put into a Pasteur pipette, with 2 L/min of continuous flow by purified and humidified air controlled by a Syntech stimulus controller (CS-55 model, Syntech, Germany). The stimuli were given at 0.5 L/min for 500ms, and the action potential signals were amplified by using a pre-amplifier (IDAC-4 USB System, Syntech, Germany) and recorded with Autospike 32 software (Syntech, Germany). The number of induced spikes were calculated by the firing spike number subtraction by spontaneous spikes number before and after 1s with the stimulus.

### Retrograde larval brain staining and confocal imaging

The 4th instar larvae were drawn into 10 uL pipette tips and fixed by the dental wax. For antennae back fillings, the fluorescent dye (Micro-Ruby, Invitrogen, Eugene, OR) was injected to the cut off antennae by the glass capillary, then the larva were maintained in a dark and humidified box at 4 °C for 12-16 h. For the whole brain imaging, the heads of larva were separated and fixed by 4% paraformaldehyde solution (PFA) for 1-2 h, then the brains was dissected in PBS buffer (1.8 mM KH2PO4, 4.3 mM Na2HPO4, 137 mM NaCl, 2.7 mM KCl, pH = 7.4) and incubating with 4% PFA at 4 ℃overnight, after that, the brains were washed with 0.5% PBST (PBS buffer containing 0.5% Triton X-100) and pre-incubated with 5% normal goat serum (NGS; Thermo Fisher Scientific, Waltham, MA, USA) at 4℃overnight. The primary antibody containing 5% NGS and 1% 3C11 (anti SYNORF1, DSHB, University of Iowa, USA) were first applied at 4℃for 3 days, after washed with 0.5% PBST, the secondary antibody 0.3% Cy2 coupled Alexa FluorTM 488 (Invitrogen, Eugene, OR) containing 1% NGS was applied at 4 ℃for 2 days, after washing with 0.5% PBST and gradient alcohol (25% to 100%), the brains were transparent by methyl salicylate, mounted on a 0.5 mm aluminum slides, and sealed with neutral balsam. The slides were excited by Cy2 and Micro-Ruby in 488nm and 546nm lasers respectively under a Laser scanning confocal microscope (LSM 980, Carl Zeiss, Jena, Germany), with 1024 × 1024 XY resolution and 1μm interval at Z-steps. ZEN blue v3.8 software (Carl Zeiss, Jena, Germany) was used to capture and visualize the larva brain and antenna back filling confocal images.

### Immunocytochemistry

Fourth instar larvae were placed on ice for 15 minutes. Their antennae were gently collected using tweezers and placed in a PBST (1.8 mM KH2PO4, 4.3 mM Na2HPO4, 137 mM NaCl, 2.7 mM KCl, pH = 7.4, with 0.5% Triton X-100) solution containing 4% PFA at 4°C for 2 days. The antennae were washed with a PBST solution for 4 × 15 minutes. Overnight block was conducted overnight in a 10% normal goat serum (NGS; Thermo Fisher Scientific, Waltham, MA, USA) blocking solution, and then incubated with the primary antibody solution at 4°C for 3 days. The primary antibody solution consisted of an anti-OR9 antibody (provided by Baijia, China) diluted 1:100 in a PBST solution containing 5% NGS. Following incubation, the antennae were serially washed with PBST 6 times for 20 minutes each and incubated with the secondary antibody 1% Goat anti-Rabbit coupled with Alexa FluorTM 568 (Invitrogen, Eugene, OR; catalog number: A-11011) for 2 days. The antennae were then washed with PBST six times, 20 minutes each and mounted in anti-Fade Medium. The negative control consisted of applying the primary antibody alone without the secondary antibody. Immunohistochemistry results were visualized using an LSM980 confocal microscope (Zeiss, Jena, Germany).

## Supplementary Figures

**Figure S1. Temporal heatmaps showing the number of *Ae. aegypti* and *Ae. albopictus* larvae detected in the ROI under food and control treatments**. Heatmap representing the number of *Ae. aegypti* and *Ae. albopictus* larvae observed in the ROI over time across four replicates for both food and control treatments. Heatmap showing the number of larvae detected inside the ROI over one hour under control and food conditions. The heatmap represents the number of larvae detected per second (1 frame per second) over the 3000-second experimental period.

**Figure S2. Maximum likelihood phylogenetic constructed using 439 chemosensory genes from *Ae. aegypti* and 479 chemosensory genes from *Ae. albopictus***. The tree includes genes from the four main chemosensory gene families: *Or, Gr, Ir, Obp & Ppk*. Bootstrap values indicated by the size of grey circles in nodes (values < 50 are not shown).

**Figure S3. Maximum likelihood phylogenetic tree of odorant receptors (*Or*) genes from *Ae. aegypti* and *Ae. albopictus***. Sequences for *Ae. aegypti* were obtained from [50], and sequences for *Ae. albopictus* were retrieved from the AalbF5 genome assembly (GCF_035046485.1) curated *OR*s. Branch lengths represent the number of substitutions per site (The scale bar refers to amino acid substitutions per site). Support levels for nodes are indicated by the size of gray circles-reflecting approximate likelihood ratio tests (Default IQ-Tree parameters). Underscore suffixes (_1) after protein names in *Ae. albopictus* refer to species-specific duplications. Letter suffixes after protein names refer to alternative transcripts of the same gene. Amino acid (AA) identity percentages between orthologs are marked in: no color (<65%), green (65-75%), yellow (75-85%), orange (85-95%) and red (95-100%) referring to the percentage of amino-acid identity between orthologs.

**Figure S4. Maximum likelihood phylogenetic tree of gustatory receptors (*Gr*) genes from *Ae. aegypti* and *Ae. albopictus***. Sequences for *Ae. aegypti* were obtained from Matthews et al. 2018 [50], and curated sequences for *Ae. albopictus* were retrieved from the AalbF5 genome assembly (GCF_035046485.1). Branch lengths represent the number of substitutions per site (The scale bar refers to amino acid substitutions per site). Support levels for nodes are indicated by the size of gray circles-reflecting approximate likelihood ratio tests (Default IQ-Tree parameters). Underscore suffixes (_1) after protein names in *Ae. albopictus* refers to species-specific duplications. Letter suffixes after protein names refer to alternative transcripts of the same gene. Amino acid (AA) identity percentages between orthologs are marked in: no color (<65%), green (65-75%), yellow (75-85%), orange (85-95%) and red (95-100%) referring to the percentage of amino-acid identity between orthologs.

**Figure S5. Maximum likelihood phylogenetic tree of ionotropic receptors (*Ir*) genes from *Ae. aegypti* and *Ae. albopictus***. Sequences for *Ae. aegypti* were obtained from Matthews et al. 2018 [50], and sequences for *Ae. albopictus* were retrieved from the AalbF5 genome assembly (GCF_035046485.1) curated *IR*s. Branch lengths represent the number of substitutions per site (The scale bar refers to amino acid substitutions per site). Support levels for nodes are indicated by the size of gray circles-reflecting approximate likelihood ratio tests (Default IQ-Tree parameters). Underscore suffixes (_1) after protein names in *Ae. albopictus* refer to species-specific duplications. Letter suffixes after protein names refer to alternative transcripts of the same gene. Amino acid (AA) identity percentages between orthologs are marked in: no color (<65%), green (65-75%), yellow (75-85%), orange (85-95%) and red (95-100%) referring to the percentage of amino-acid identity between orthologs.

**Figure S6. Maximum likelihood phylogenetic tree of odorant-binding proteins (*Obp*) genes from *Ae. aegypti* and *Ae. albopictus***. Sequences for *Ae. aegypti* were obtained from Matthews et al. 2018 [50], and curated sequences for *Ae. albopictus* were retrieved from the AalbF5 genome assembly (GCF_035046485.1). Branch lengths represent the number of substitutions per site (The scale bar refers to amino acid substitutions per site). Support levels for nodes are indicated by the size of gray circles-reflecting approximate likelihood ratio tests (default IQ-Tree parameters). Underscore suffixes (_1) after protein names in *Ae. albopictus* refer to species-specific duplications. Letter suffixes after protein names refer to alternative transcripts of the same gene. Amino acid (AA) identity percentages between orthologs are marked in: no color (<65%), green (65-75%), yellow (75-85%), orange (85-95%) and red (95-100%) referring to the percentage of amino-acid identity between orthologs.

**Figure S7. Maximum likelihood phylogenetic tree of pick-pocket receptors proteins (*Ppk*) genes from *Ae. aegypti* and *Ae. albopictus***. Sequences for *Ae. aegypti* were obtained from Matthews et al. 2018 [50], and sequences for *Ae. albopictus* were retrieved from the AalbF5 genome assembly (GCF_035046485.1). Branch lengths represent the number of substitutions per site (The scale bar refers to amino acid substitutions per site). Support levels for nodes are indicated by the size of gray circles-reflecting approximate likelihood ratio tests (default IQ-Tree parameters). Letter suffixes after protein names refer to alternative transcripts of the same gene. Amino acid (AA) identity percentages between orthologs are marked in: no color (<65%), green (65-75%), yellow (75-85%), orange (85-95%) and red (95-100%) referring to the percentage of amino-acid identity between orthologs.

**Figure S8. Developmental *OR* expression in *Ae. aegypti* and *Ae. albopictus***. Heatmap representation of *Or* expression levels in larval and adult antennae in each species. Log_2_ fold-change values for each gene are indicated by circles (black circles, *Ae. albopictus*; brown circles, *Ae. aegypti*). Solid circles represent differentially expressed genes (*p* < 0.001), while the lack of significant gene expression differences are marked with open circles. Expression values are based on mean normalized reads (MRN) from three biological replicates.

**Figure S9. Developmental *Gr* expression in *Ae. aegypti* and *Ae. albopictus***. Heatmap representation of *Gr* expression levels in larval and adult antennae in each species. Log_2_ fold-change values for each gene are indicated by circles (black circles, *Ae. albopictus*; brown circles, *Ae. aegypti*). Solid circles represent differentially expressed genes (*p* < 0.001), while the lack of significant gene expression differences are marked with open circles. Expression values are based on mean normalized reads (MRN) from three biological replicates.

**Figure S10. Developmental *Ir* expression in *Ae. aegypti* and *Ae. albopictus***. Heatmap representation of *Ir* expression levels in larval and adult antennae in each species. Log_2_ fold-change values for each gene are indicated by circles (black circles, *Ae. albopictus*; brown circles, *Ae. aegypti*). Solid circles represent differentially expressed genes (*p* < 0.001), while the lack of significant gene expression differences are marked with open circles. Expression values are based on mean normalized reads (MRN) from three biological replicates.

**Figure S11. Developmental *OBP* expression in *Ae. aegypti* and *Ae. albopictus***. Heatmap representation of *Obp* expression levels in larval and adult antennae in each species. Log_2_ fold-change values for each gene are indicated by circles (black circles, *Ae. albopictus*; brown circles, *Ae. aegypti*). Solid circles represent differentially expressed genes (*p* < 0.001), while the lack of significant gene expression differences are marked with open circles. Expression values are based on mean normalized reads (MRN) from three biological replicates.

**Figure S12. Developmental *Ppk* expression in *Ae. aegypti* and *Ae. albopictus***. Heatmap representation of *Ppk* expression levels in larval and adult antennae in each species. Log_2_ fold-change values for each gene are indicated by circles (black circles, *Ae. albopictus*; brown circles, *Ae. aegypti*). Solid circles represent differentially expressed genes (*P* < 0.001), while the lack of significant gene expression differences are marked with open circles. Expression values are based on mean normalized reads (MRN) from three biological replicates.**Figure S13**. Differential expression Scatter plots for *Ae. aegypti* and *Ae. albopictus. Or, Gr, Ir, Obp* and *Ppk* plots are ordered from top to bottom. Genes with significant stage differential expression (Padj < 0.001) are shown in turquoise or blue for larva/adult biased respectively, while non-significant genes between the stages are depicted in gray.

**Figure S14. Single-sensillum recording of the larval cone. A**) Ventral view of the larval head, the reference electrode inserted into the eye and the recording electrode placed above the antenna. **B**) Representative traces showing responses to 1-octen-3-ol (positive control) and increasing doses of indole and skatole. *Ae. albopictus* traces are on the right while *Ae. aegypti* are on the left.

**Figure S15. Oocyte traces of AalbOR2, 9 & 10 to indole and skatole in black and red respectively**. Raw traces of all replicates and summarizing the table found in the supplementary folder (DOI 10.5281/zenodo.15603830 /Pharmacology folder).

**Figure S16. Replicates of anti-*Or9* in-situ immunolocalizations in the *Ae. albopictus* larval antenna**.

**Figure S17. *Ae. albopictus* replicates of anterograde brain staining using left and right antennal dye injections**. Solid triangles indicate contralateral projections.

**Figure S18. *Ae. aegypti* replicates of anterograde brain staining using left and right antennal dye injections**. Solid triangles indicate contralateral projections.

## Author Contributions

JDB conceived the study. GC and RMW conducted the gene annnottations and orthology analyses of *Ae. albopictus* genes. DP and EY collected the antennal tissues used for RNA-seq. DP and ESS designed and conducted the behavioral experiments. DP analyzed conducted the bioinformatic analyses the supervision FK, PAP and JDB. HK conducted the RT-qPCR, larval immuno-histochemistry, larva SSR and larval brain anterograde staining in *Ae. albopictus* under the supervision of YW. ZY conducted the RT-qPCR, larval immuno-histochemistry, larva SSR and larval brain anterograde staining in *Ae. aegypti* under the supervision of YW. JDB, DP, YW and PAP designed the figures. JB wrote the manuscript with individual contributions from all the authors.

## Acknowledgments

We would like to express our gratitude to Yuri Vainer, Michal Arbel, Doron Zaada, and Yael Arien for their valuable contributions. This research was supported by a grant from the Israel Science Foundation (719/21) to JDB. This research was also supported by grants from the Ministry of Science & Technology, Israel to PAP (grant agreement numbers 3-16795 and 3-17985). Funding was also provided by the German-Israeli Middle East Project Cooperation of the German Research Foundation (SCHE 1833/7-1 to PAP).

## Competing Interest Statement

The authors declare no conflict of interests.

## Notes

https://doi.org/10.5281/zenodo.17292855

### Competing Interest Statement

The authors have declared no competing interest.

### Summary of Updates

In this revision, we have added new Aedes aegypti data, including qRT-PCR gene expression, sensory recordings, and brain imaging, thereby completing the interspecies comparison. We also included links to several supplementary materials, such as Ae. albopictus F5 chemosensory gene annotations, transcriptomic metadata, raw behavioral videos, real-time gene expression data, and pharmacological receptor recordings.

## References

1. M. U. G. Kraemer, M. E. Sinka, K. A. Duda, A. Q. Mylne, F. M. Shearer, C. M. Barker, C. G. Moore, R. G. Carvalho, G. E. Coelho, W. Van Bortel, G. Hendrickx, F. Schaffner, I. R. F. Elyazar, H.-J. Teng, O. J. Brady, J. P. Messina, D. M. Pigott, T. W. Scott, D. L. Smith, G. R. W. Wint, S. I. Hay, The global distribution of the arbovirus vectors Aedes aegypti and Aedes albopictus. eLife 4, e08347 (2015). 10.7554/eLife.08347

2. C. M. Gossner, E. Ducheyne, F. Schaffner, Increased risk for autochthonous vector-borne infections transmitted by Aedes albopictus in continental Europe. Euro Surveill. 23, 1800268 (2018). 10.2807/1560-7917.ES.2018.23.24.1800268

3. L. P. Lounibos, L. D. Kramer, Invasiveness of Aedes aegypti and Aedes albopictus and vectorial capacity for Chikungunya virus. J. Infect. Dis. 214 (Suppl 5), S453–S458 (2016). 10.1093/infdis/jiw285

4. M. Bonizzoni, G. Gasperi, X. Chen, A. A. James, The invasive mosquito species Aedes albopictus: current knowledge and future perspectives. Trends Parasitol. 29, 460–468 (2013). 10.1016/j.pt.2013.07.003

5. J. Soghigian, A. Gloria-Soria, V. Robert, G. Le Goff, A.-B. Failloux, J. R. Powell, Genetic evidence for the origin of Aedes aegypti, the yellow fever mosquito, in the southwestern Indian Ocean. Mol. Ecol. 29, 3593–3606 (2020). 10.1111/mec.15590

6. J. Soghigian, C. Sither, S. A. Justi, et al., Phylogenomics reveals the history of host use in mosquitoes. Nat. Commun. 14, 6252 (2023). 10.1038/s41467-023-41764-y

7. J. R. Powell, W. J. Tabachnick, History of domestication and spread of Aedes aegypti—a review. Mem. Inst. Oswaldo Cruz 108 (Suppl 1), 11–17 (2013). 10.1590/0074-0276130395

8. M. U. G. Kraemer, R. C. Reiner Jr., O. J. Brady, J. P. Messina, M. Gilbert, D. M. Pigott, D. Yi, K. Johnson, L. Earl, L. B. Marczak, S. Shirude, N. D. Weaver, D. Bisanzio, T. A. Perkins, S. Lai, X. Lu, P. Jones, G. E. Coelho, R.G. Carvalho, W. Van Bortel, N. Golding, Past and future spread of the arbovirus vectors Aedes aegypti and Aedes albopictus. Nat. Microbiol. 4, 854–863 (2019). 10.1038/s41564-019-0376-y

9. L. Mousson, C. Dauga, T. Garrigues, F. Schaffner, M. Vazeille, A.-B. Failloux, Phylogeography of Aedes aegypti and Aedes albopictus (Diptera: Culicidae) based on mitochondrial DNA variations. Genet. Res. 86, 1–11 (2005). 10.1017/S0016672305007627

10. S. K. Panigrahi, et al., Review of feeding, biting, and resting behavior of Aedes aegypti and Aedes albopictus. Int. J. Mosq. Res. 11, 1–11 (2024). 10.22271/23487941.2024.v11.i3a.771

11. L. P. Lounibos, et al., Asymmetric evolution of photoperiodic diapause in temperate and tropical invasive populations of Aedes albopictus (Diptera: Culicidae). Ann. Entomol. Soc. Am. 96, 512–518 (2003). 10.1603/0013-8746(2003)096[0512:AEOPDI]2.0.CO;2

12. A. Kolimenakis, et al., The role of urbanisation in the spread of Aedes mosquitoes and the diseases they transmit—A systematic review. PLoS Negl. Trop. Dis. 15, e0009631 (2021). 10.1371/journal.pntd.0009631

13. M. Braks, et al., Convergent habitat segregation of Aedes aegypti and Aedes albopictus (Diptera: Culicidae) in southeastern Brazil and Florida. J. Med. Entomol. 40, 785–794 (2003). 10.1603/0022-2585-40.6.785

14. S. A. Juliano, L. P. Lounibos, Ecology of invasive mosquitoes: effects on resident species and on human health. Ecol. Lett. 8, 558–574 (2005). 10.1111/j.1461-0248.2005.00755.x

15. P. Nie, J. Feng, Niche and range shifts of Aedes aegypti and Aedes albopictus suggest that the latecomer shows a greater invasiveness. Insects 14, 810 (2023). 10.3390/insects14100810

16. J. Zhou, S. Liu, H. Liu, et al., Interspecific mating bias may drive Aedes albopictus displacement of Aedes aegypti during its range expansion. PNAS Nexus 1, pgac041 (2022). 10.1093/pnasnexus/pgac041

17. S. A. Brennan, I. C. Grob, C. E. Bartz, J. K. Baker, Y. Jiang, Displacement of Aedes albopictus by Aedes aegypti in Gainesville, Florida. J. Am. Mosq. Control Assoc. 37, 93–97 (2021). 10.2987/20-6992.1

18. G. F. O’Meara, L. F. Evans Jr., A. D. Gettman, J. P. Cuda, Spread of Aedes albopictus and decline of Aedes aegypti (Diptera: Culicidae) in Florida. J. Med. Entomol. 32, 554–562 (1995). 10.1093/jmedent/32.4.554

19. R. S. Nasci, S. G. Hare, F. S. Willis, Interspecific mating between Louisiana strains of Aedes albopictus and Aedes aegypti in the field and laboratory. J. Am. Mosq. Control Assoc. 5, 416–421 (1989).

20. B. Kamgang, T. A. Wilson-Bahun, H. Irving, et al., Geographical distribution of Aedes aegypti and Aedes albopictus (Diptera: Culicidae) and genetic diversity of invading populations of Aedes albopictus in the Republic of the Congo. Wellcome Open Res. 3, 79 (2018). 10.12688/wellcomeopenres.14659.3

21. L. Bagny Beilhe, H. Delatte, S. A. Juliano, D. Fontenille, S. Quilici, Ecological interactions in Aedes species on Reunion Island. Med. Vet. Entomol. 27, 387–397 (2013). 10.1111/j.1365-2915.2012.01062.x

22. M. A. H. Braks, N. A. Honório, L. P. Lounibos, R. Lourenço-De-Oliveira, S. A. Juliano, Interspecific competition between two invasive species of container mosquitoes, Aedes aegypti and Aedes albopictus (Diptera: Culicidae), in Brazil. Ann. Entomol. Soc. Am. 97, 130–139 (2004). 10.1603/0013-8746(2004)097[0130:ICBTIS]2.0.CO;2

23. R. G. Carvalho, R. Lourenço-de-Oliveira, I. A. Braga, Updating the geographical distribution and frequency of Aedes albopictus in Brazil with remarks regarding its range in the Americas. Mem. Inst. Oswaldo Cruz 109, 787– 796 (2014). 10.1590/0074-0276140304

24. J. H. Hobbs, E. A. Hughes, B. H. Eichold 2nd, Replacement of Aedes aegypti by Aedes albopictus in Mobile, Alabama. J. Am. Mosq. Control Assoc. 7, 488–489 (1991).

25. S. K. Gilotra, L. E. Rozeboom, N. C. Bhattacharya, Observations on possible competitive displacement between populations of Aedes aegypti Linnaeus and Aedes albopictus Skuse in Calcutta. Bull. World Health Organ. 37, 437– 446 (1967).

26. L. Bagny, H. Delatte, N. Elissa, S. Quilici, D. Fontenille, Aedes (Diptera: Culicidae) vectors of arboviruses in Mayotte (Indian Ocean): distribution area and larval habitats. J. Med. Entomol. 46, 198–207 (2009). 10.1603/033.046.0204

27. L. Kaplan, D. Kendell, D. Robertson, et al., Aedes aegypti and Aedes albopictus in Bermuda: extinction, invasion, invasion and extinction. Biol. Invasions 12, 3277–3288 (2010). 10.1007/s10530-010-9721-z

28. C. Caminade, M. I. Vasquez, H. Herodotou, G. Notarides, C. Pavlou, F. Fayad, A. Papakonstantinou, M. Violaris, D. Petric, W. Mamai, A. M. Tompkins, J. Bouyer, The invasions of Aedes aegypti and Aedes albopictus in Cyprus: Current situation, risk modelling, and public health implications for the wider Eastern Mediterranean region. bioRxiv [Preprint] (2024). 10.1101/2024.12.17.628941

29. S. A. Juliano, Species introduction and replacement among mosquitoes: interspecific resource competition or apparent competition? Ecology 79, 255–268 (1998). 10.1890/0012-9658(1998)079[0255:SIARAM]2.0.CO;2

30. D. A. Yee, B. Kesavaraju, S. A. Juliano, Interspecific differences in feeding behavior and survival under food-limited conditions for larval Aedes albopictus and Aedes aegypti (Diptera: Culicidae). Ann. Entomol. Soc. Am. 97, 720–728 (2004). 10.1603/0013-8746(2004)097[0720:IDIFBA]2.0.CO;2

31. J. J. Skiff, D. A. Yee, Behavioral differences among four co-occurring species of container mosquito larvae: Effects of depth and resource environments. J. Med. Entomol. 51, 375–381 (2014). 10.1603/ME13159

32. L. Houri-Zeevi, M. M. Walker, J. Razzauti, A. Sharma, H. A. Pasolli, L. B. Vosshall, Mosquito sex under lock and key. bioRxiv [Preprint] (2025). 10.1101/2025.04.11.648401

33. M. N. Andersson, C. I. Keeling, R. F. Mitchell, Genomic content of chemosensory genes correlates with host range in wood-boring beetles (Dendroctonus ponderosae, Agrilus planipennis, and Anoplophora glabripennis). BMC Genomics 20, 690 (2019). 10.1186/s12864-019-6054-x

34. E. K. Lutz, C. Lahondère, C. Vinauger, J. A. Riffell, Olfactory learning and chemical ecology of olfaction in disease vector mosquitoes: a life history perspective. Curr. Opin. Insect Sci. 20, 75–83 (2017). 10.1016/j.cois.2017.03.002

35. R. Y. Zacharuk, S. G. Blue, Ultrastructure of the peg and hair sensilla on the antenna of larval Aedes aegypti (L.). J. Morphol. 135, 433–455 (1973). 10.1002/jmor.1051350403

36. R. Y. Zacharuk, L. R. Yin, S. G. Blue, Fine structure of the antenna and its sensory cone in larvae of Aedes aegypti (L.). J. Morphol. 135, 273–297 (1971). 10.1002/jmor.1051350303

37. F. Scolari, et al., Imaging and spectral analysis of autofluorescence distribution in larval head structures of mosquito vectors. Eur. J. Histochem. 66, 3462 (2022). 10.4081/ejh.2022.3462

38. S. B. McIver, Sensilla of mosquitoes (Diptera: Culicidae). J. Med. Entomol. 19, 489–535 (1982). 10.1093/jmedent/19.5.489

39. M. Bui, J. Shyong, E. K. Lutz, et al., Live calcium imaging of Aedes aegypti neuronal tissues reveals differential importance of chemosensory systems for life-history-specific foraging strategies. BMC Neurosci. 20, 27 (2019). 10.1186/s12868-019-0511-y

40. J. Bohbot, et al., Molecular characterization of the Aedes aegypti odorant receptor gene family. Insect Mol. Biol. 16, 525–537 (2007). 10.1111/j.1365-2583.2007.00748.x

41. Y. Xia, G. Wang, D. Buscariollo, R. J. Pitts, H. Wenger, L. J. Zwiebel, The molecular and cellular basis of olfactory-driven behavior in Anopheles gambiae larvae. Proc. Natl. Acad. Sci. U.S.A. 105, 6433–6438 (2008). 10.1073/pnas.0801007105

42. B. J. Matthews, C. S. McBride, M. DeGennaro, et al., The neurotranscriptome of the Aedes aegypti mosquito. BMC Genomics 17, 32 (2016). 10.1186/s12864-015-2239-0

43. F. Lombardo, M. Salvemini, C. Fiorillo, T. Nolan, L. J. Zwiebel, J. M. Ribeiro, B. Arcà, Deciphering the olfactory repertoire of the tiger mosquito Aedes albopictus. BMC Genomics 18, 770 (2017). 10.1186/s12864-017-4144-1

44. W. S. Leal, Odorant reception in insects: roles of receptors, binding proteins, and degrading enzymes. Annu. Rev. Entomol. 58, 373–391 (2013). 10.1146/annurev-ento-120811-153635

45. J. D. Bohbot, P. L. Jones, G. Wang, R. J. Pitts, G. M. Pask, L. J. Zwiebel, Conservation of indole-responsive odorant receptors in mosquitoes reveals an ancient olfactory trait. Chem. Senses 36, 149–160 (2011). 10.1093/chemse/bjq105

46. J. D. Bohbot, J. C. Dickens, Characterization of an enantioselective odorant receptor in the yellow fever mosquito Aedes aegypti. PLoS One 4, e7032 (2009). 10.1371/journal.pone.0007032

47. D. Ruel, et al., Supersensitive odorant receptor underscores pleiotropic roles of indoles in mosquito ecology. Front. Cell. Neurosci. 12, 533 (2018). 10.3389/fncel.2018.00533

48. J. D. Bohbot, J. C. Dickens, Insect repellents: modulators of mosquito odorant receptor activity. PLoS One 5, e12138 (2010). 10.1371/journal.pone.0012138

49. S. Shankar, C. J. McMeniman, An updated antennal lobe atlas for the yellow fever mosquito Aedes aegypti. PLoS Negl. Trop. Dis. 14, e0008729 (2020). 10.1371/journal.pntd.0008729

50. B. J. Matthews, O. Dudchenko, S. B. Kingan, et al., Improved reference genome of Aedes aegypti informs arbovirus vector control. Nature 563, 501–507 (2018). 10.1038/s41586-018-0692-z

51. A. Sánchez-Gracia, F. G. Vieira, J. Rozas, Molecular evolution of the major chemosensory gene families in insects. Heredity 103, 208–216 (2009). 10.1038/hdy.2009.55

52. C. J. McMeniman, et al., Multimodal integration of carbon dioxide and other sensory cues drives mosquito attraction to humans. Cell 156, 1060–1071 (2014). 10.1016/j.cell.2013.12.044

53. D. Ma, et al., Structural basis for sugar perception by Drosophila gustatory receptors. Science 383 (6685), eadj2609 (2024). 10.1126/science.adj2609

54. S.Y. Zhao, P.L. Wu, J.Y. Fu, Y.M. Wu, H.K. Liu, L.J. Cai, J.B. Gu, X.H. Zhou, X.G. Chen, Gustatory receptor 11 is involved in detecting the oviposition water of the Asian tiger mosquito, Aedes albopictus. Parasites Vectors 17, 367 (2024). 10.1186/s13071-024-06452-w

55. J. I. Raji, C. J. Potter, Chemosensory ionotropic receptors in human host-seeking mosquitoes. Curr. Opin. Insect Sci. 54, 100967 (2022). 10.1016/j.cois.2022.100967

56. E. K. Lutz, K. T. Ha, J. A. Riffell, Distinct navigation behaviors in Aedes, Anopheles, and Culex mosquito larvae. J. Exp. Biol. 223 (Pt 7), jeb221218 (2020). 10.1242/jeb.221218

57. L. Weisskopf, S. Schulz, P. Garbeva, Microbial volatile organic compounds in intra-kingdom and inter-kingdom interactions. Nat. Rev. Microbiol. 19, 391–404 (2021). 10.1038/s41579-020-00508-1

58. K. E. Levy Karin, M. Mirdita, J. Söding, MetaEuk—sensitive, high-throughput gene discovery and annotation for large-scale eukaryotic metagenomics. Microbiome 8, 48 (2020). 10.1186/s40168-020-00808-x

59. D. Kuznetsov, F. Tegenfeldt, M. Manni, M. Seppey, M. Berkeley, E. V. Kriventseva, E. M. Zdobnov, OrthoDB v11: annotation of orthologs in the widest sampling of organismal diversity. Nucleic Acids Res. 51 (D1), D445–D451 (2023). 10.1093/nar/gkac998

60. K. Katoh, J. Rozewicki, K. D. Yamada, MAFFT online service: Multiple sequence alignment, interactive sequence choice, and visualization. Brief Bioinform. 20, 1160–1166 (2018). 10.1093/bib/bbx108

61. S. Kuraku, C. M. Zmasek, O. Nishimura, K. Katoh, aLeaves facilitates on-demand exploration of metazoan gene family trees on MAFFT sequence alignment server with enhanced interactivity. Nucleic Acids Res. 41 (Web Server issue), W22–W28 (2013). 10.1093/nar/gkt389

62. L. T. Nguyen, H. A. Schmidt, A. Von Haeseler, B. Q. Minh, IQ-TREE: A fast and effective stochastic algorithm for estimating maximum-likelihood phylogenies. Mol. Biol. Evol. 32, 268–274 (2015). 10.1093/molbev/msu300

63. A. M. Bolger, M. Lohse, B. Usadel, Trimmomatic: A flexible trimmer for Illumina sequence data. Bioinformatics 30, 2114–2120 (2014). 10.1093/bioinformatics/btu170

64. D. Kim, B. Langmead, S. L. Salzberg, HISAT: A fast spliced aligner with low memory requirements. Nat. Methods 12, 357–360 (2015). 10.1038/nmeth.3317

65. Y. Liao, G. K. Smyth, W. Shi, FeatureCounts: An efficient general-purpose program for assigning sequence reads to genomic features. Bioinformatics 30, 923–930 (2014). 10.1093/bioinformatics/btt656

66. M. I. Love, W. Huber, S. Anders, Moderated estimation of fold change and dispersion for RNA-seq data with DESeq2. Genome Biol. 15, 550 (2014). 10.1186/s13059-014-0550-8

