## Supplementary figures and images for "Chemosensory mechanisms of larval foraging and competitive advantage in the invasive mosquito *Aedes albopictus*"

### Figure S1

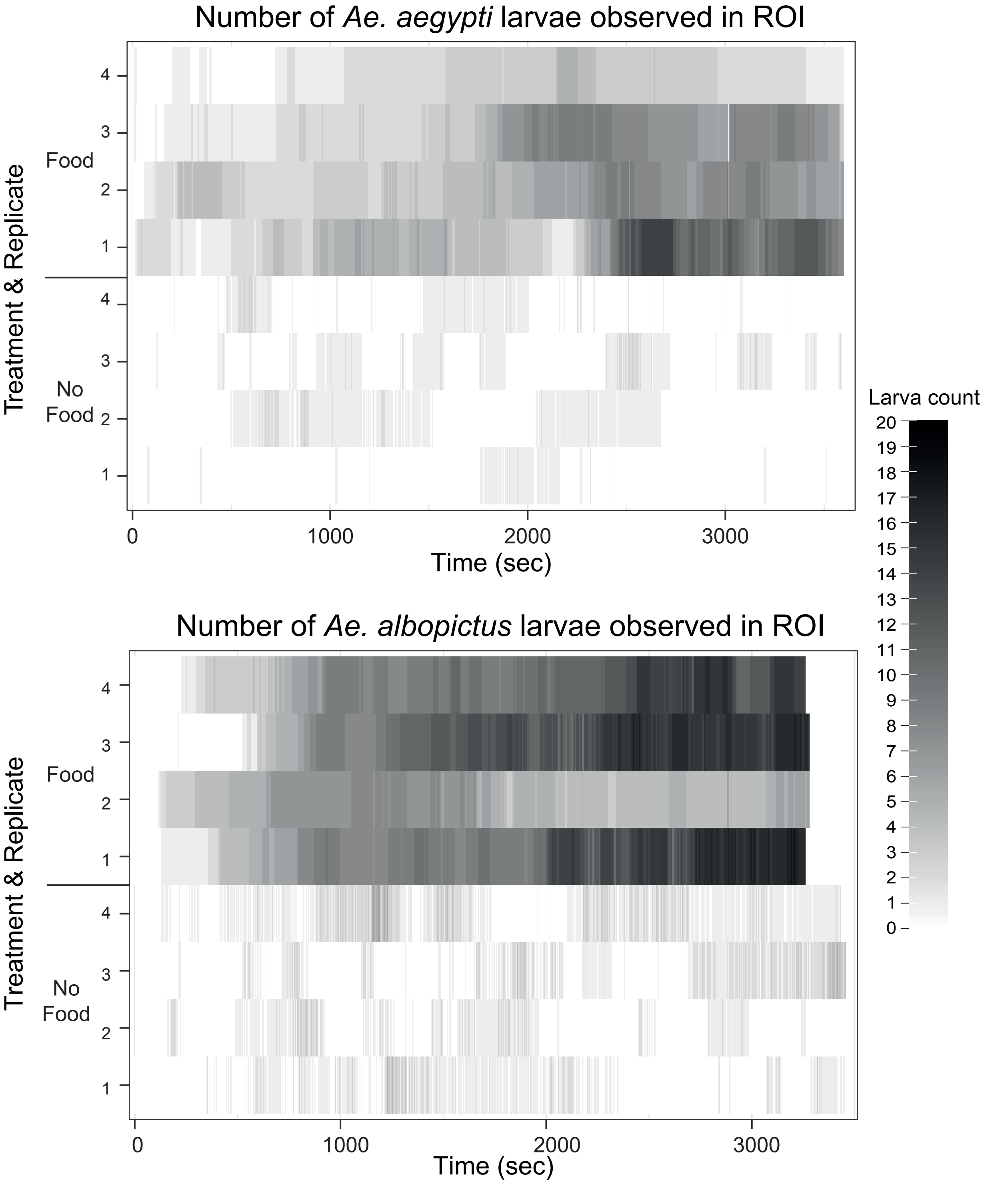

### Figure S2

Figure S2

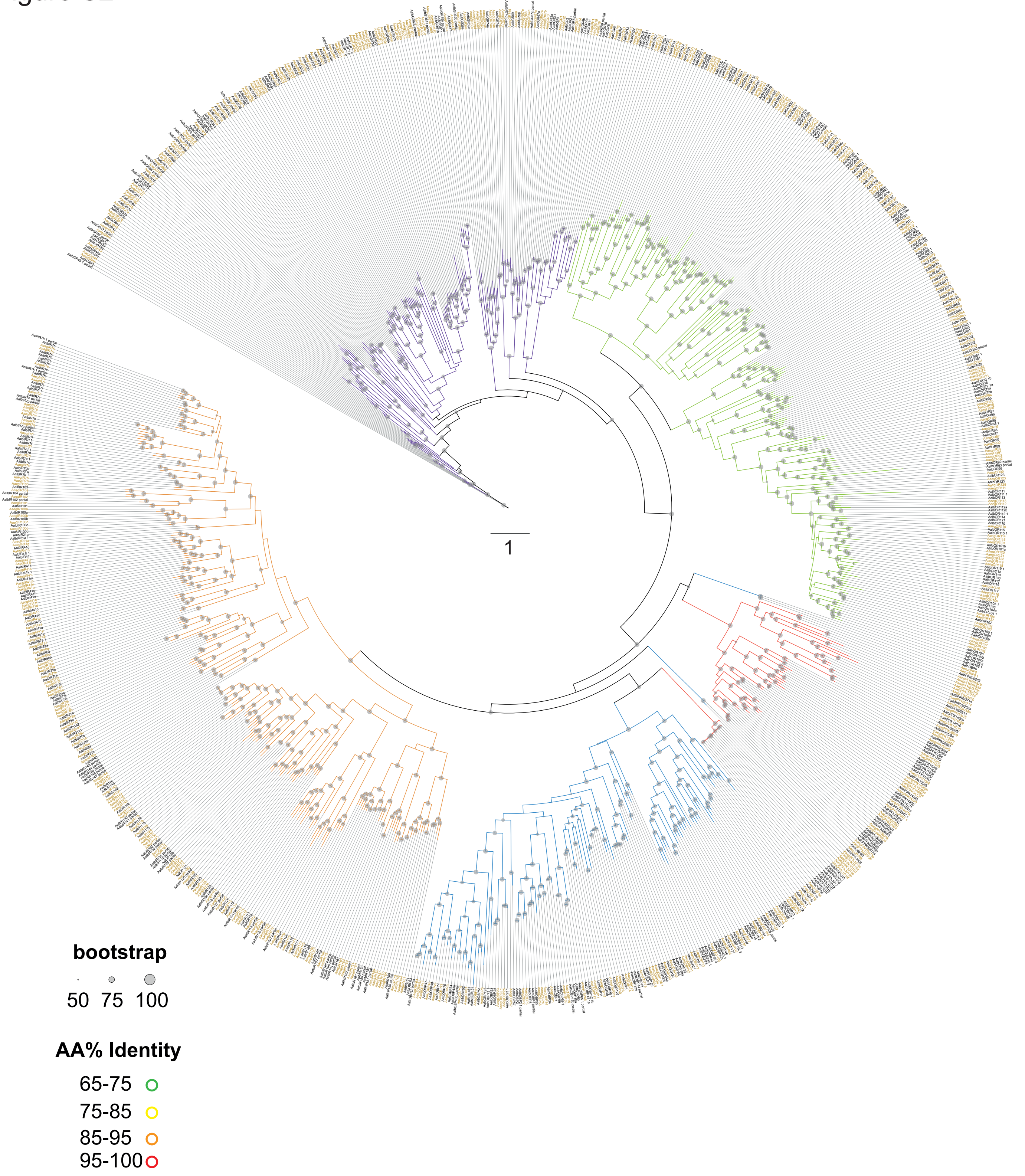

### Figure S3

Figure S3

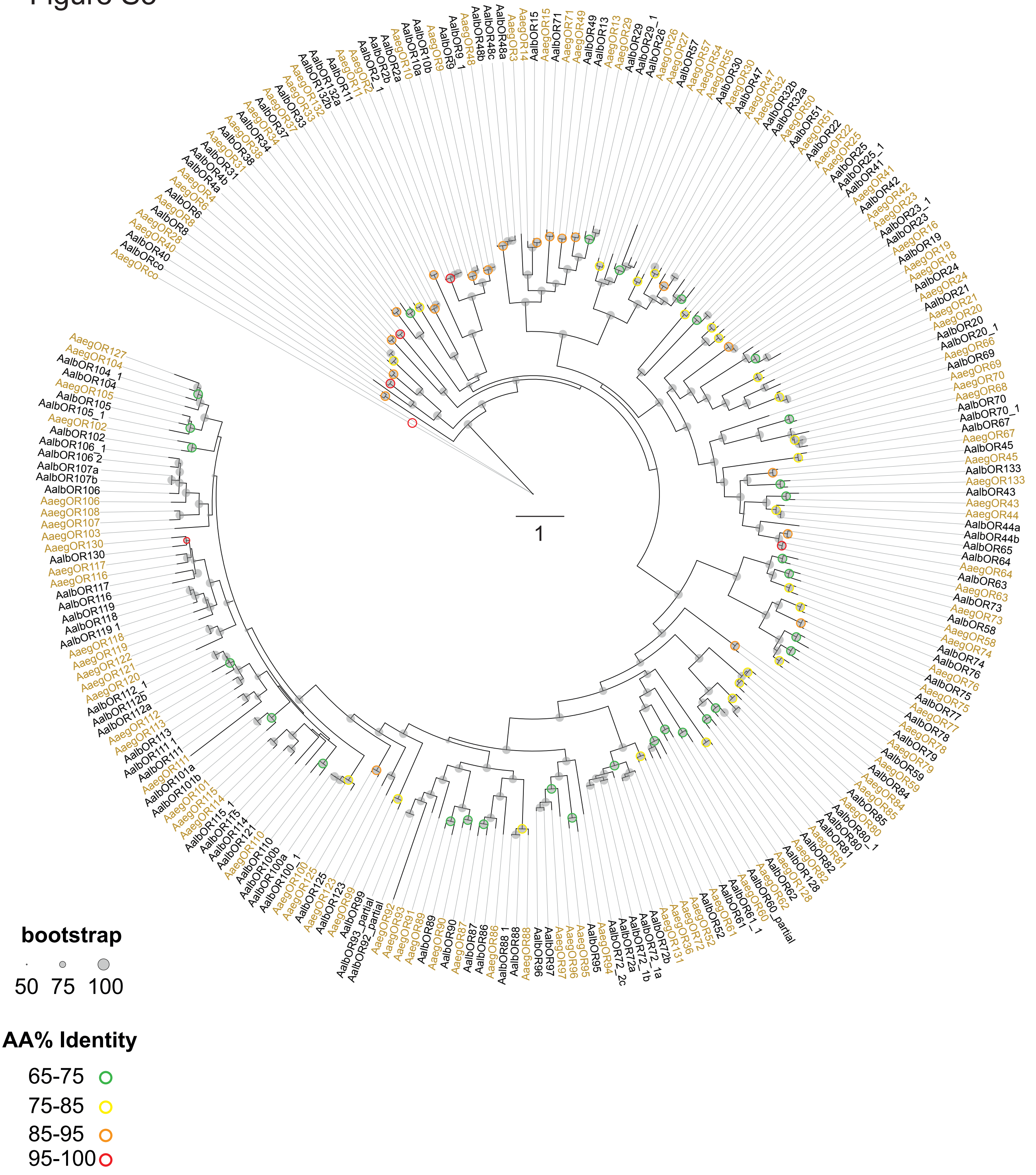

### Figure S4

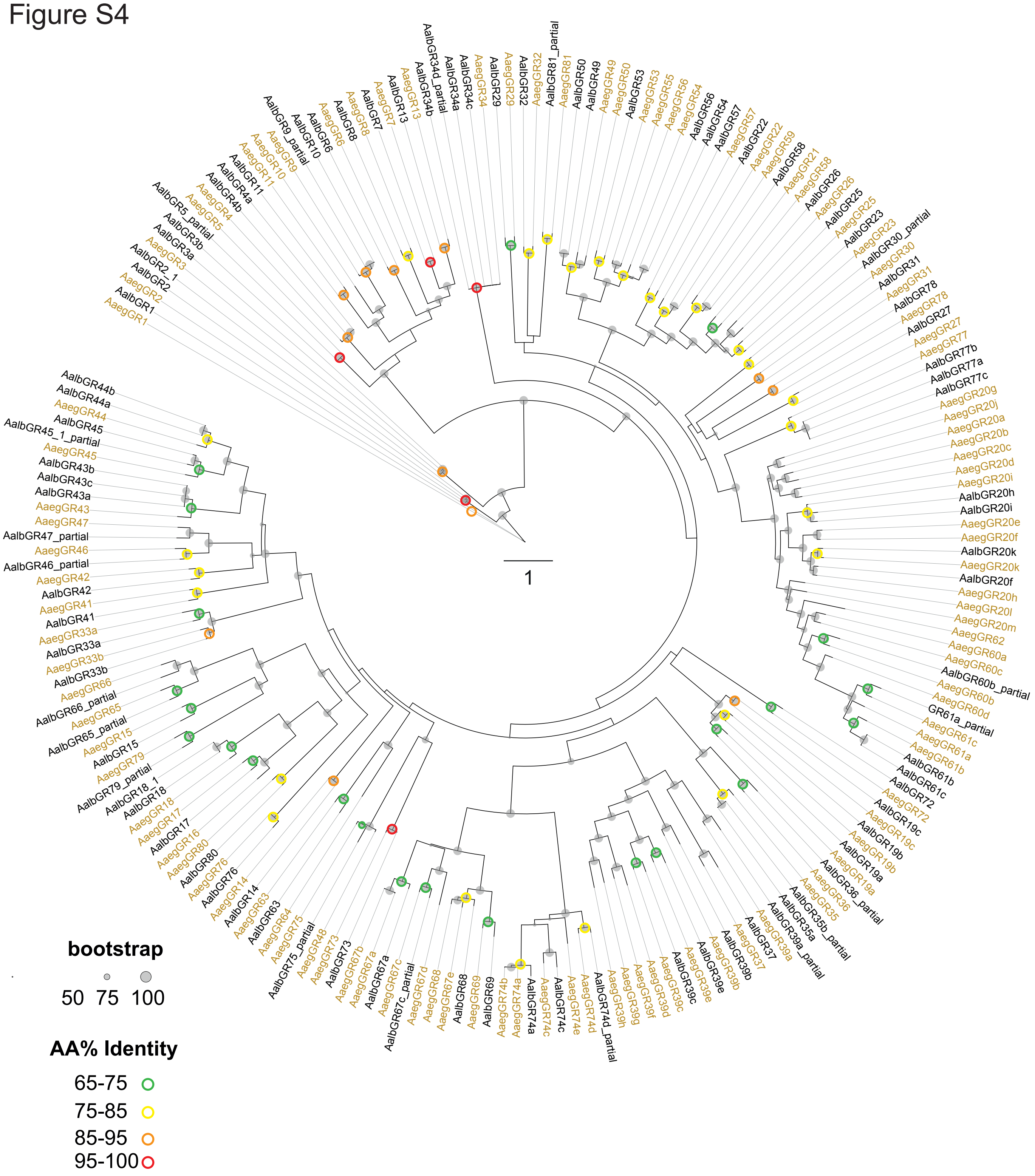

### Figure S5

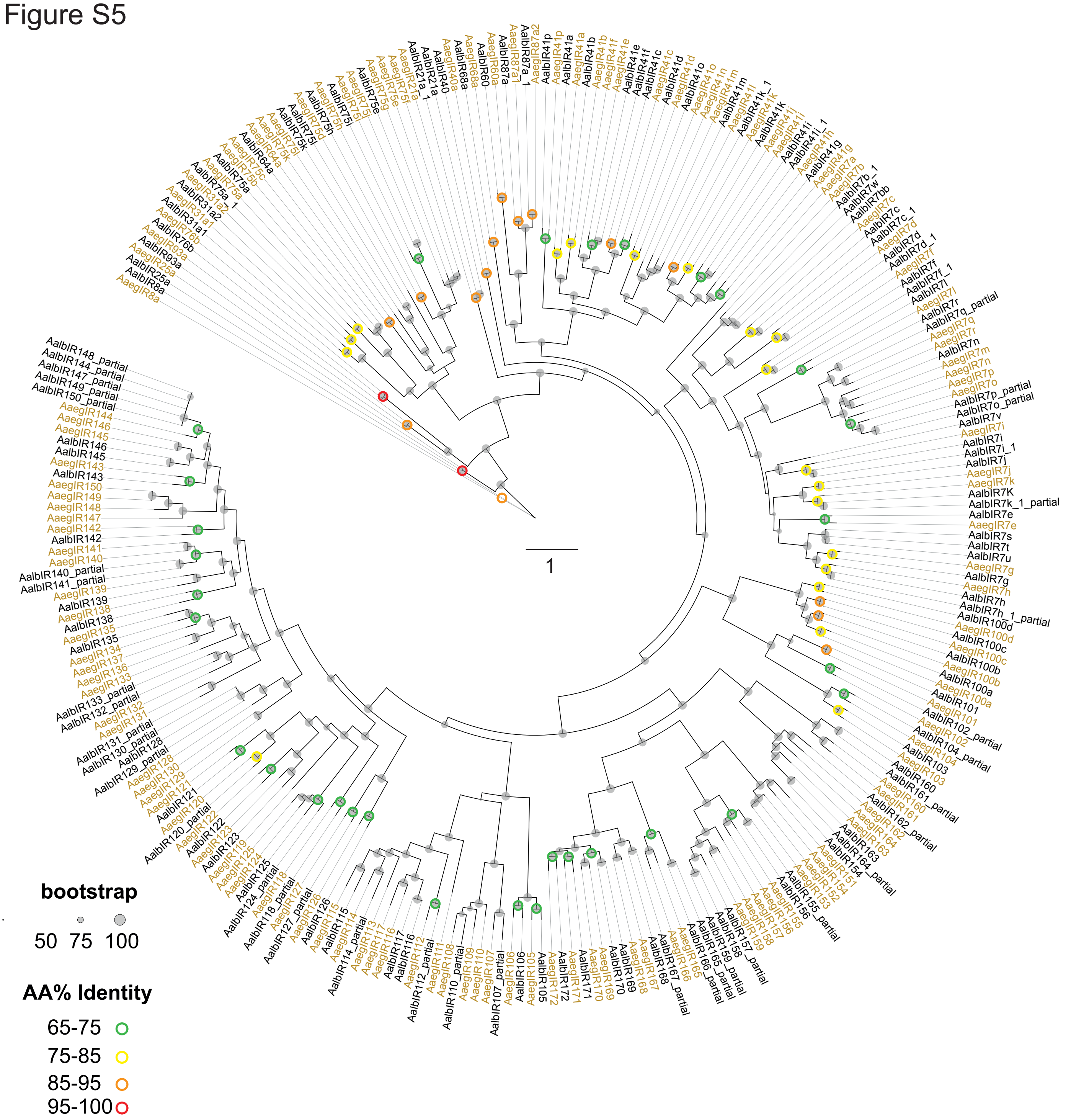

### Figure S6

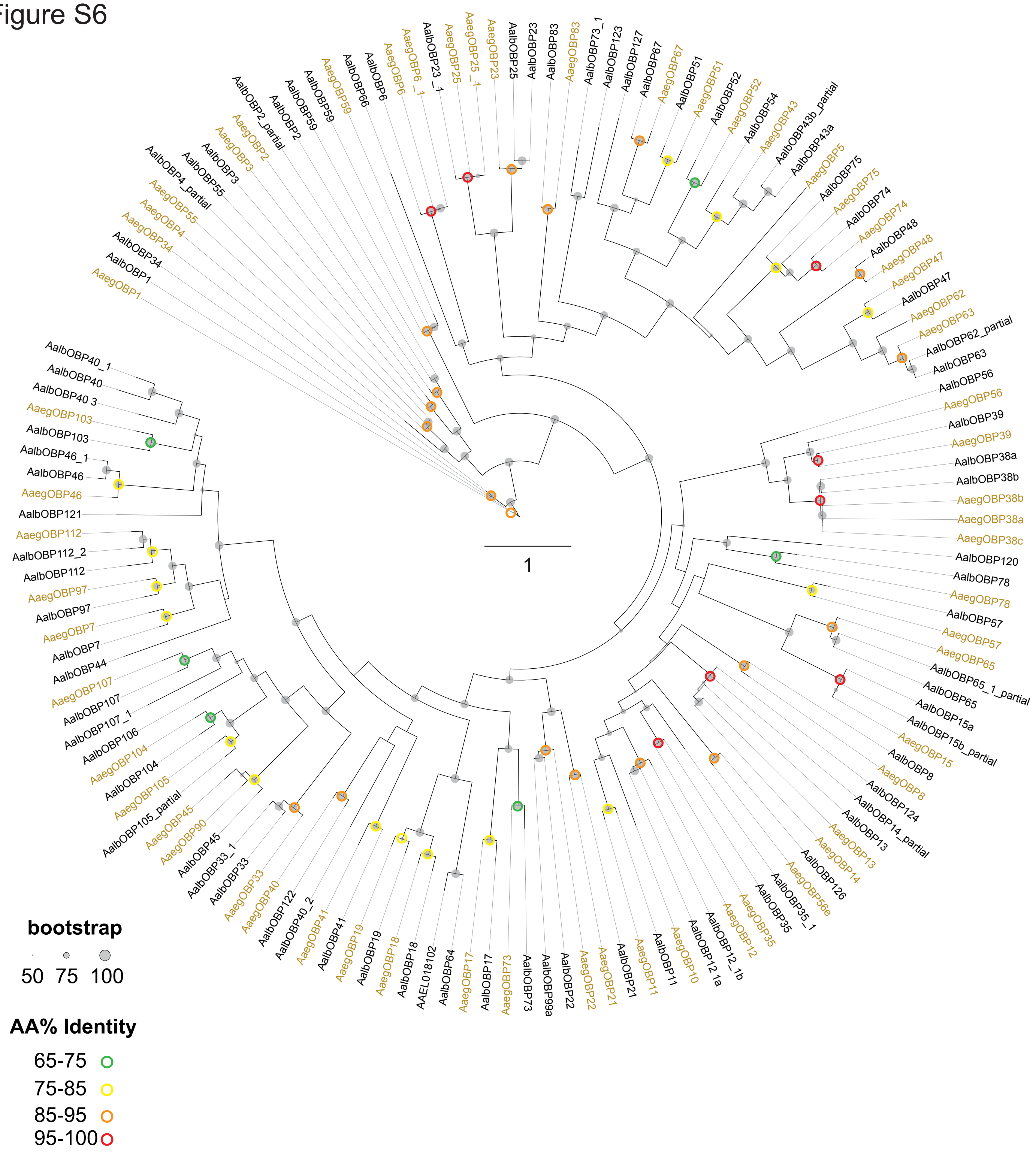

### Figure S7

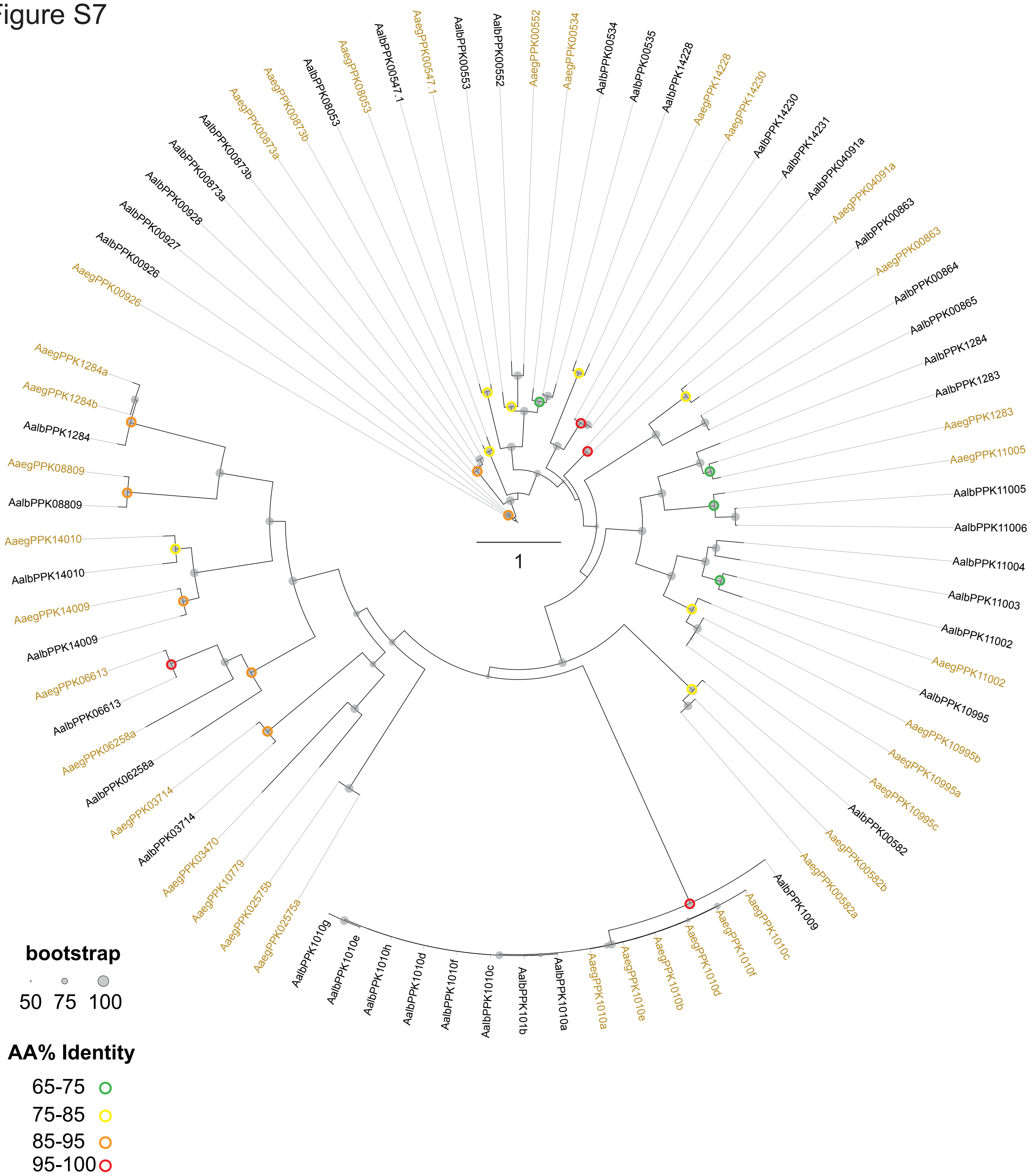

### Figure S8

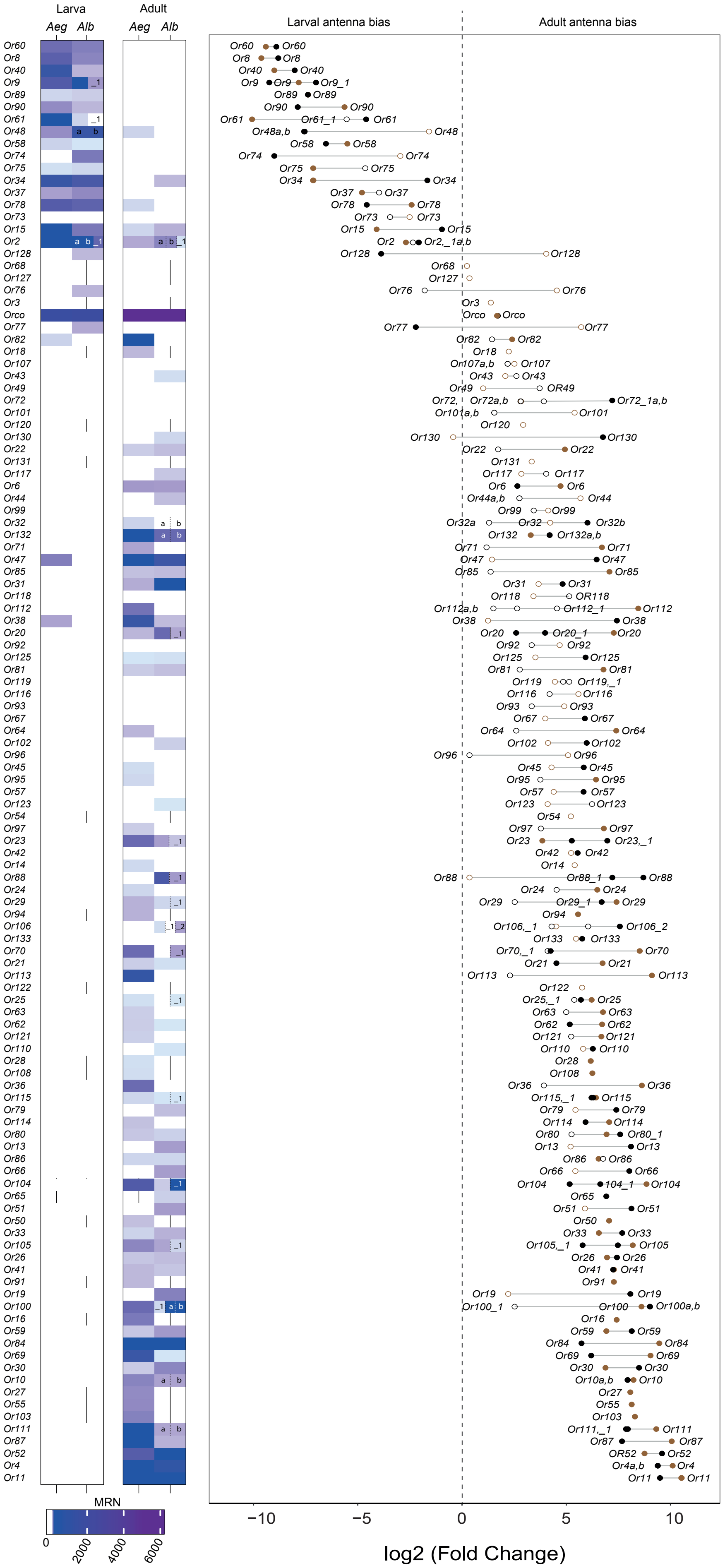

### Figure S9

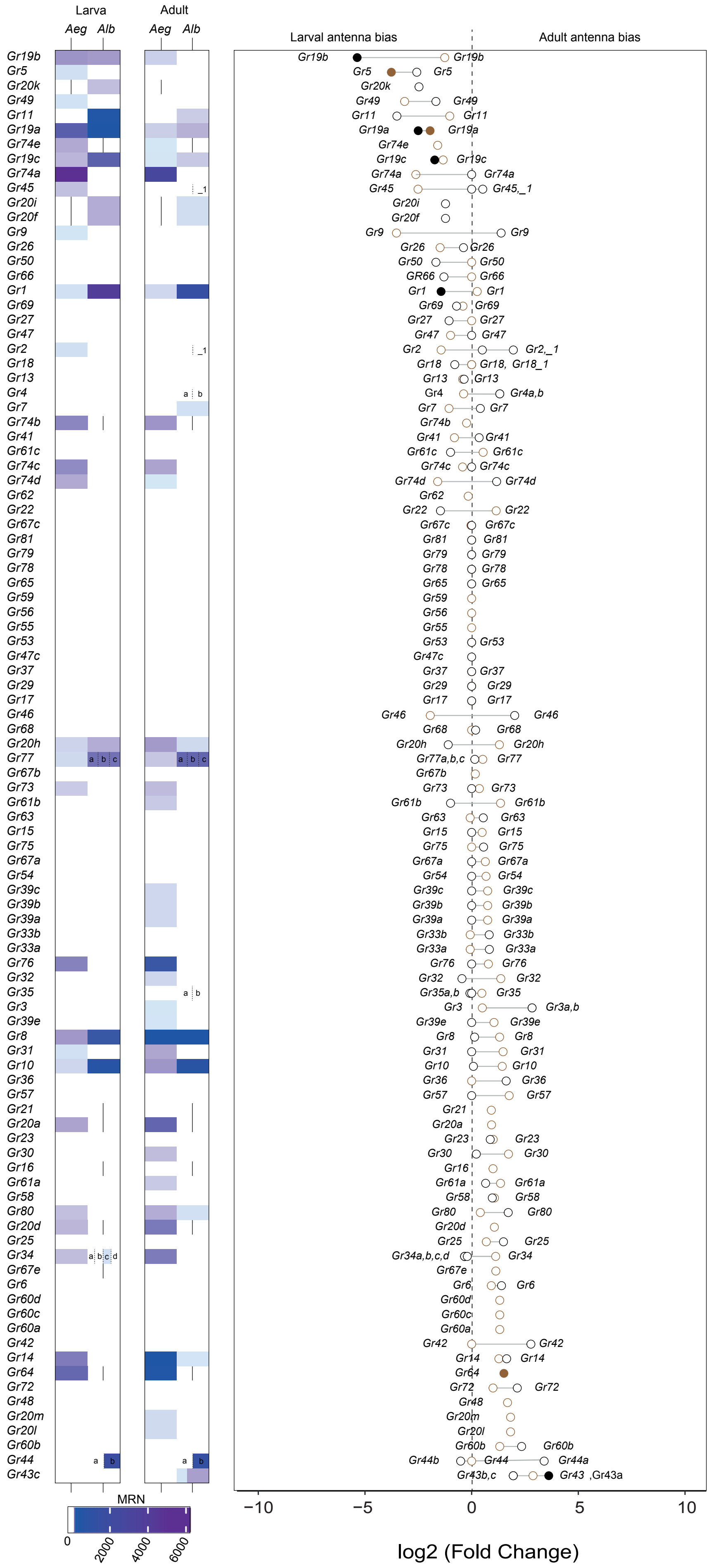

### Figure S10

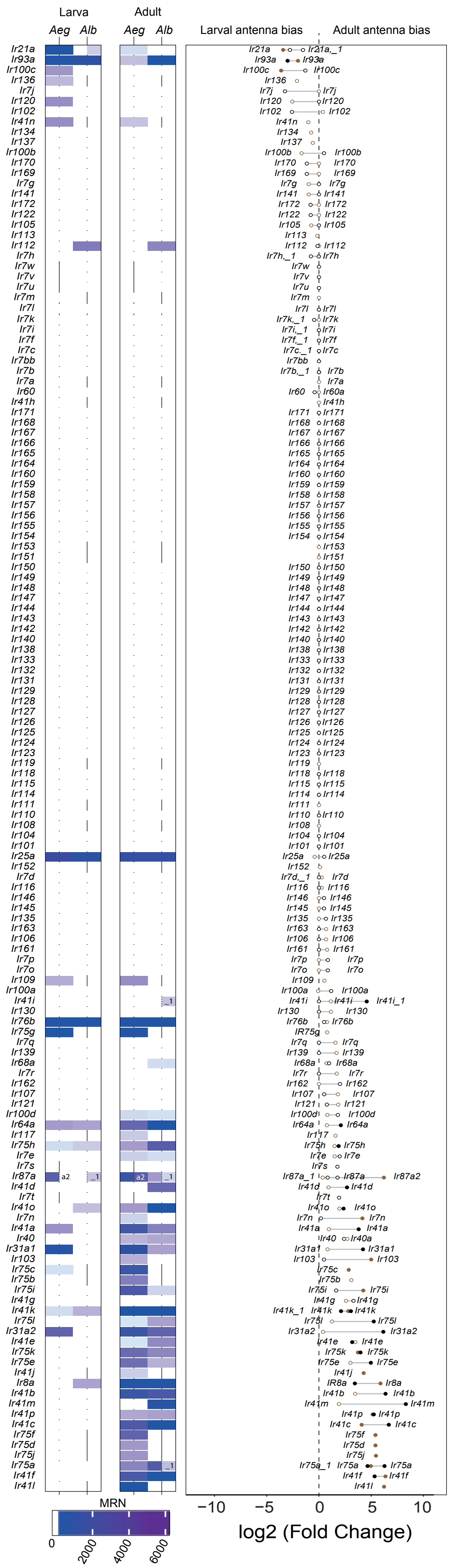

### Figure S11

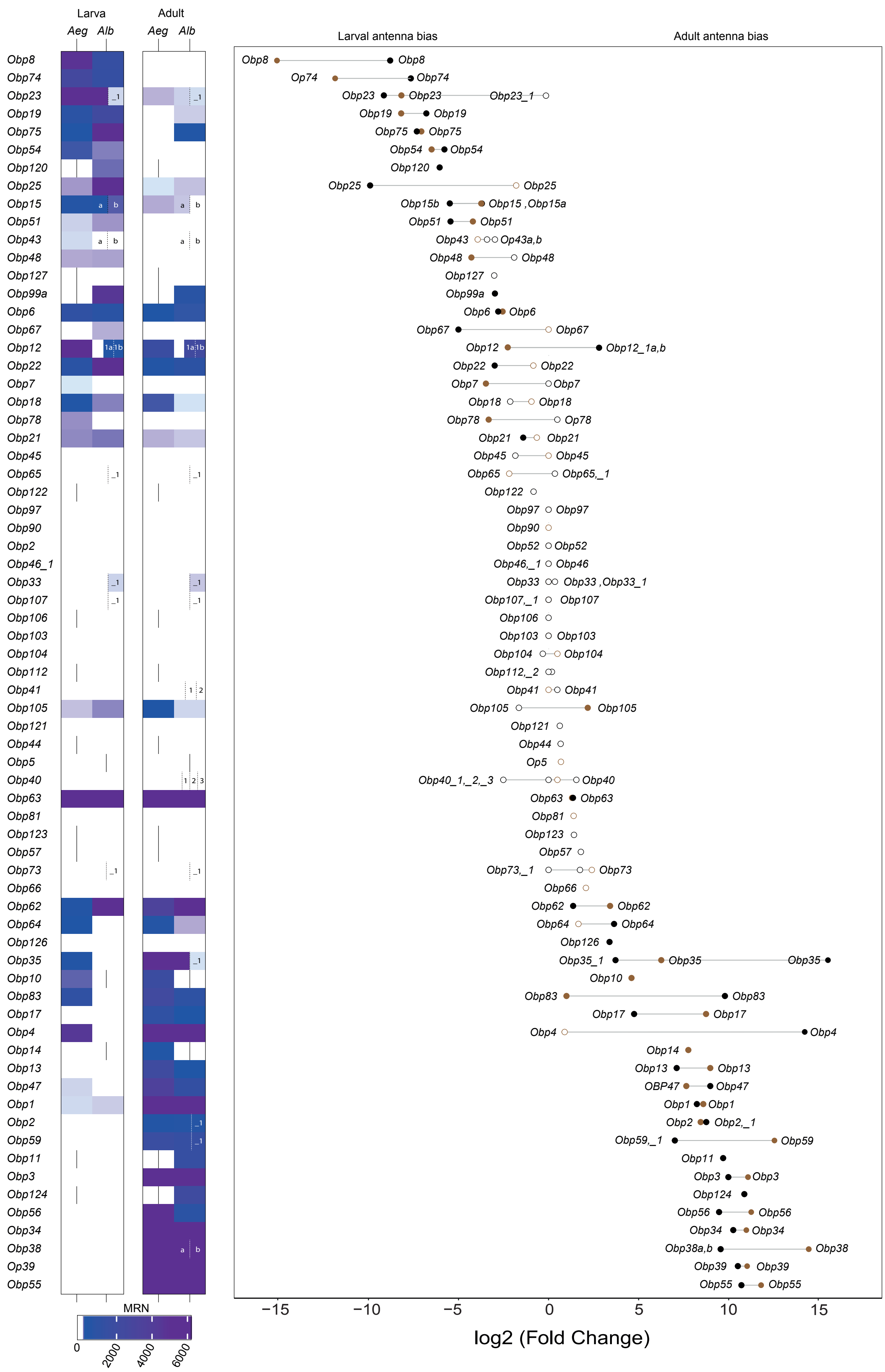

### Figure S12

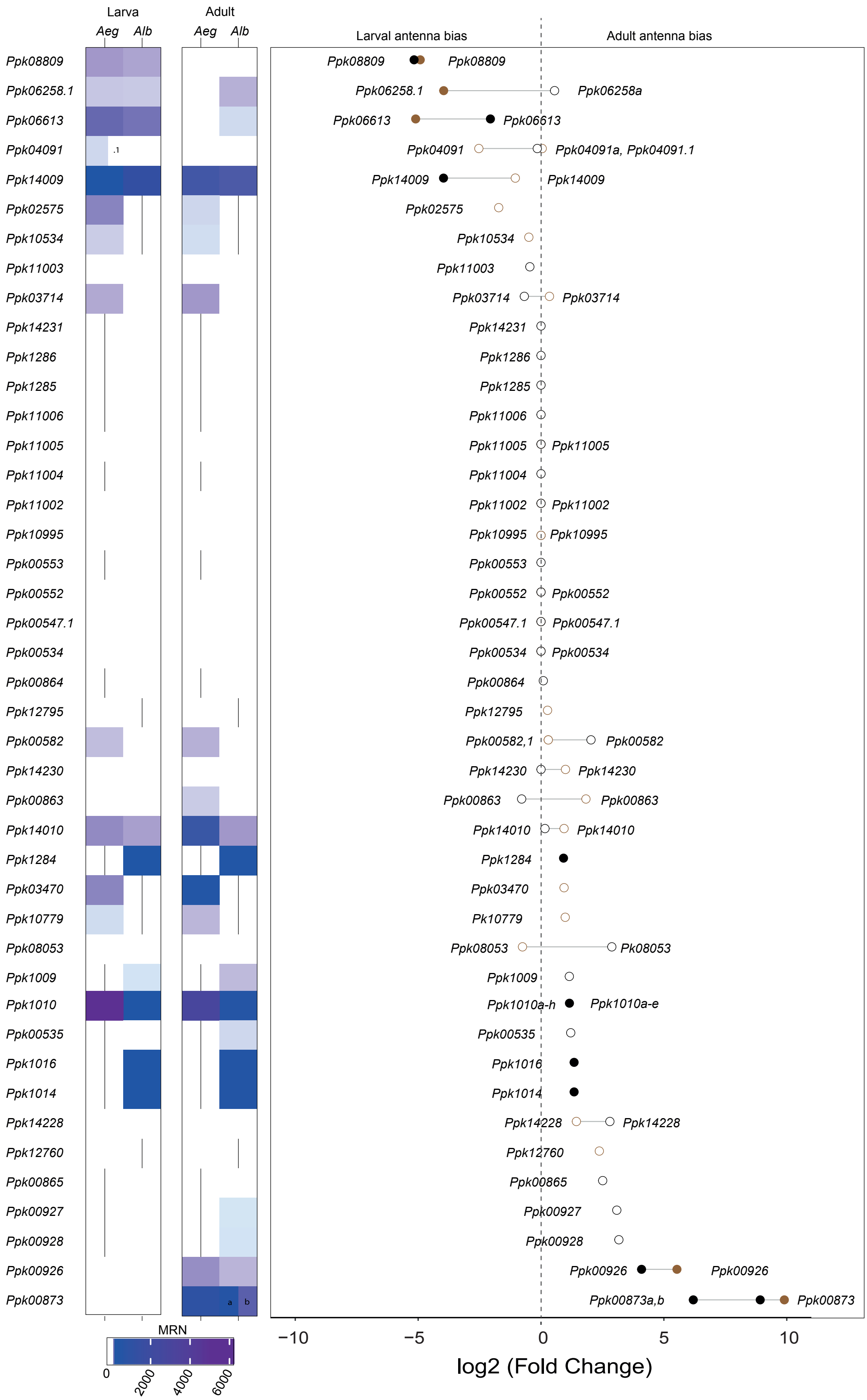

### Figure S13

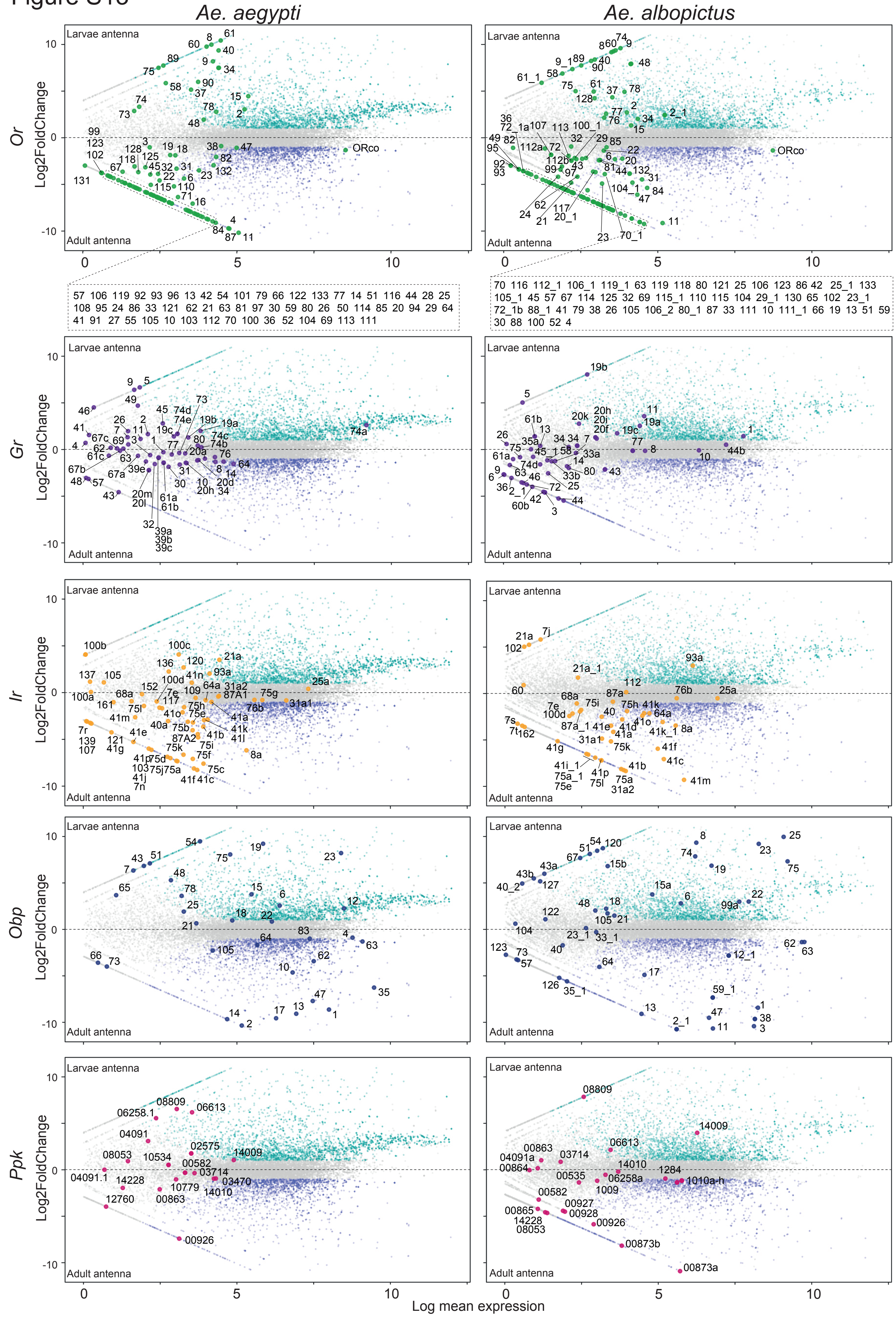

### Figure S14

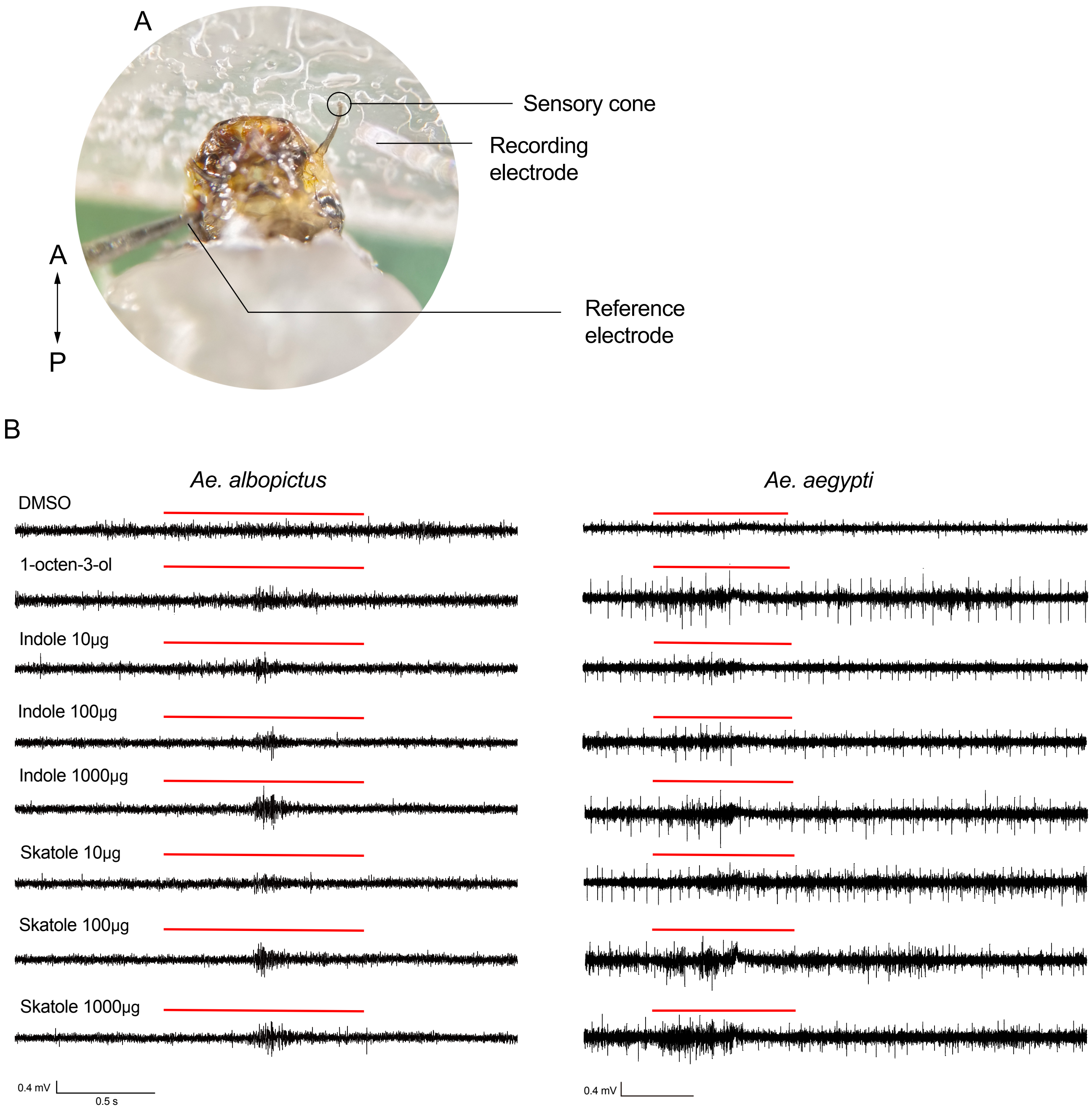

### Figure S15

Figure S15

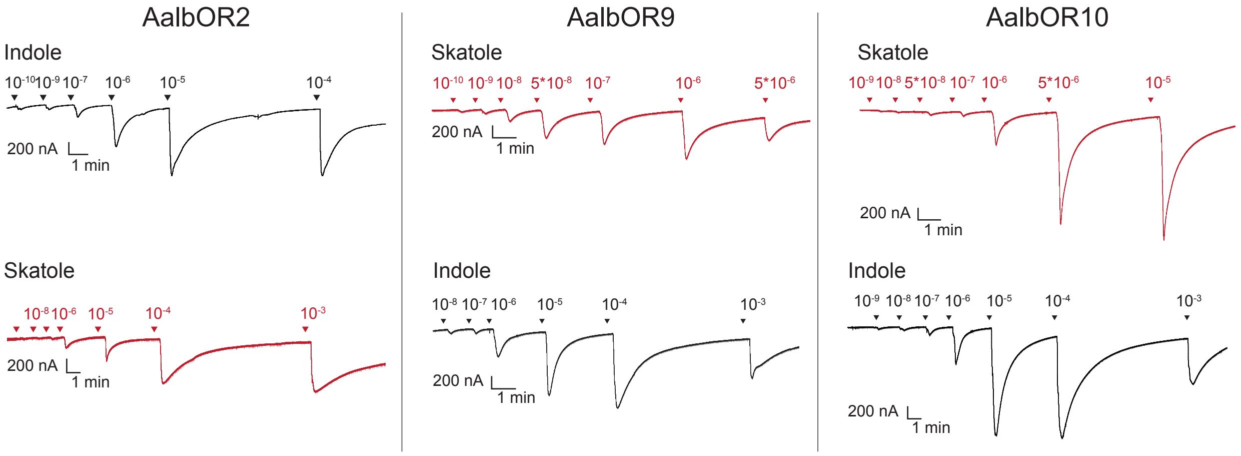


**bootstrap**

50 75 100

**AA% Identity**

65-75
75-85
85-95
95-100

### Figure S16

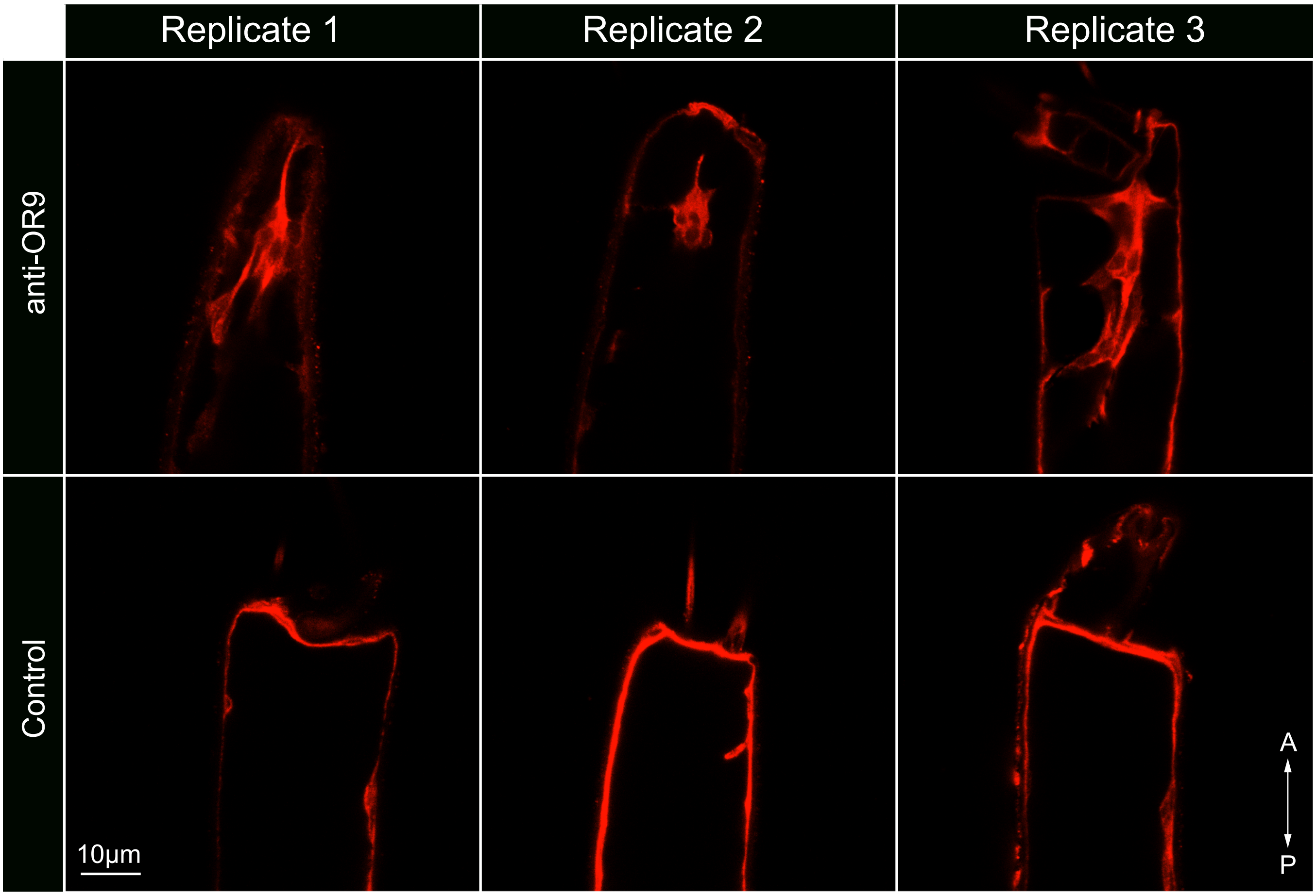

### Figure S17

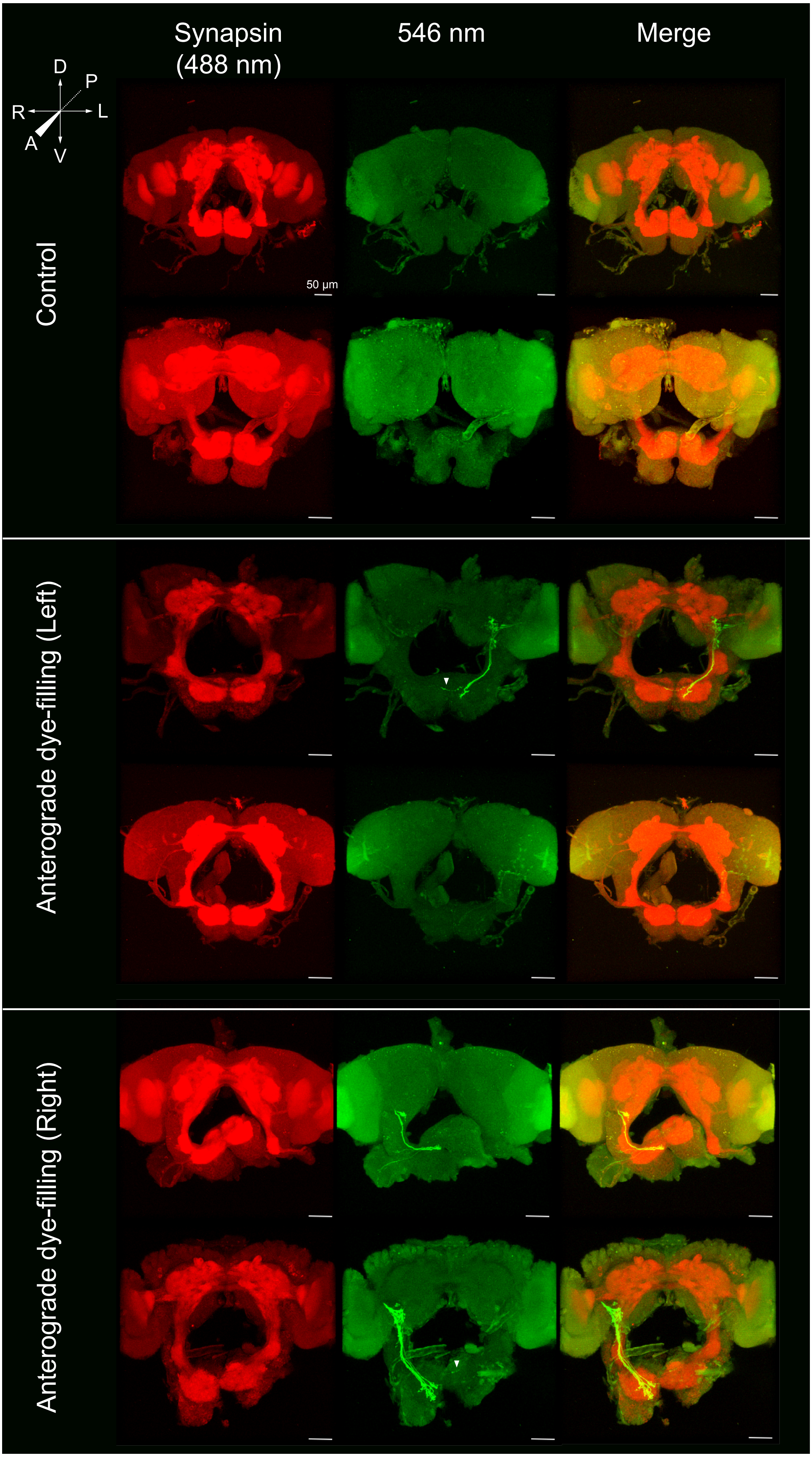

### Figure S18

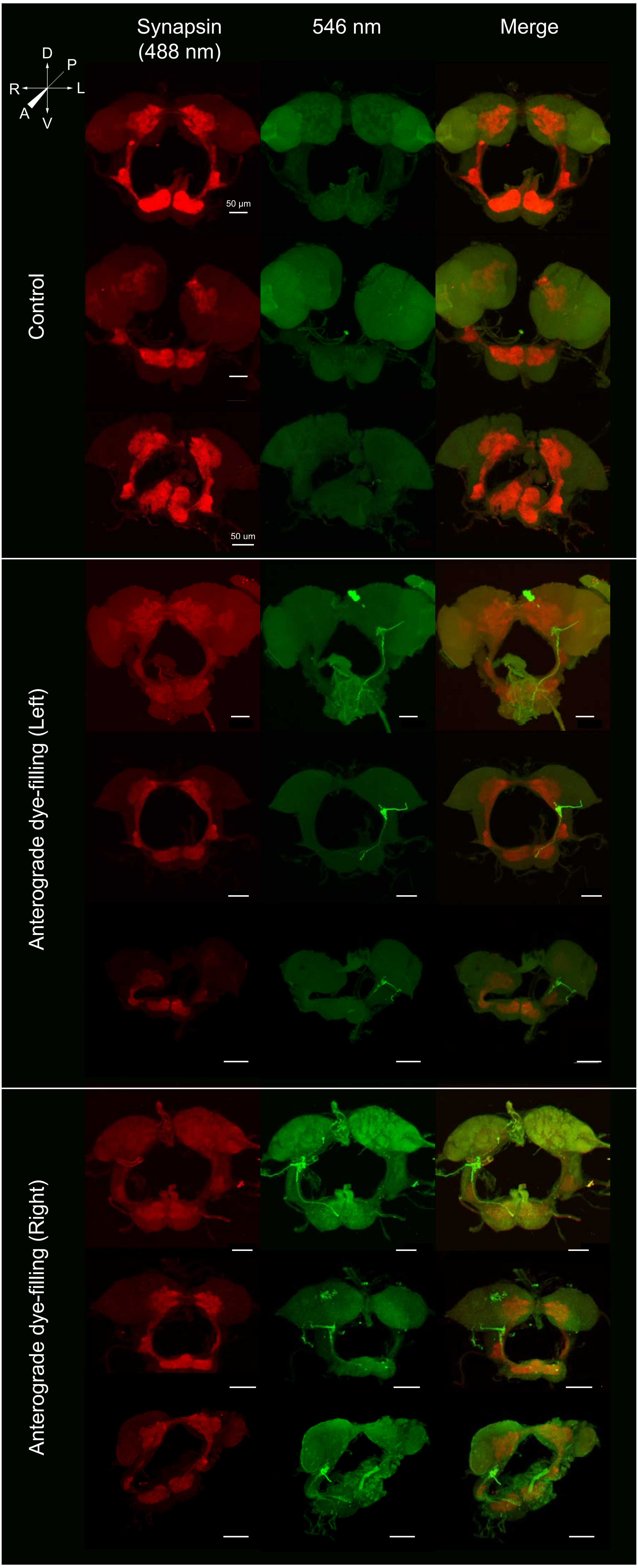
